# Unexpected versatility in the metabolism and ecophysiology of globally relevant nitrite-oxidizing *Nitrotoga* bacteria

**DOI:** 10.1101/317552

**Authors:** Andrew M. Boddicker, Annika C. Mosier

## Abstract

Nitrite-oxidizing bacteria (NOB) play a critical role in the mitigation of nitrogen pollution from freshwater systems by metabolizing nitrite to nitrate, which is removed via assimilation, denitrification, or anammox. Recent studies revealed that NOB are phylogenetically and metabolically diverse, yet most of our knowledge of NOB comes from only a few cultured representatives. Using enrichment methods and genomic sequencing, we identified four novel *Candidatus* Nitrotoga NOB species from freshwater sediments and water column samples in Colorado, USA. Genome assembly revealed highly conserved 16S rRNA gene sequences, but a surprisingly broad metabolic potential including genes for nitrogen, sulfur, hydrogen, and organic carbon metabolism. Genomic predictions suggest that *Nitrotoga* can metabolize in low oxygen or anaerobic conditions, which may support a previously unrecognized environmental niche. An array of antibiotic and metal resistance genes likely allows *Nitrotoga* to withstand environmental pressures in impacted systems. Phylogenetic analyses reveal a deeply divergent nitrite oxidoreductase alpha subunit (NxrA) not represented in any other NOB, suggesting a novel evolutionary trajectory for *Nitrotoga*. *Nitrotoga*-like 16S rRNA gene sequences were prevalent in globally distributed environments. This work considerably expands our knowledge of the *Nitrotoga* genus and improves our understanding of their role in the global nitrogen cycle.

## INTRODUCTION

Increasing anthropogenic sources of nitrogen have led to environmental risks including eutrophication, increased greenhouse gas emissions, and acidification. Nitrite-oxidizing bacteria (NOB) play a critical role in mitigating the harmful effects of nitrogen pollution by linking nitrogen sources to nitrogen removal. Specifically, nitrite pools from natural (e.g., ammonia oxidation) or anthropogenic (e.g., fertilizer) sources are converted into nitrate, which is assimilated or removed as inert nitrogen gas via denitrification or anammox pathways. Thus, understanding the diversity, metabolism, and environmental limits of NOB is important for controlling and managing elevated nitrogen in contaminated ecosystems.

Despite the environmental relevance of NOB, the group is understudied in part due to slow growth, taking as long as 12 years for isolation (Lebedeva, *et al*. 2008). Assiduous cultivation efforts (Alawi *et al*., 2007; Daims *et al*., 2015; Sorokin *et al*., 2012; van Kessel *et al*., 2015; Watson *et al*., 1986; Watson & Waterbury, 1971), as well as single-cell (Ngugi *et al*., 2015) and metagenomic (Pinto *et al*., 2015) sequencing studies are beginning to illuminate the diversity of nitrite oxidizers. The known NOB belong to four phyla and seven different genera, three of which were discovered within the last decade: *Candidatus* Nitrotoga, *Nitrolancea*, and *Candidatus* Nitromaritima (Alawi *et al*., 2007; Ngugi *et al*., 2015; Sorokin *et al*., 2012). NOB are physiologically versatile, utilizing nitrite oxidation, complete ammonia oxidation (comammox within the *Nitrospira*) (Daims *et al*., 2015; van Kessel *et al*., 2015), as well as organic carbon, hydrogen, and sulfur metabolisms to drive growth (Koch *et al*., 2016; Daims *et al*., 2016; Füssel *et al*., 2017).

*Nitrotoga* is an understudied genus of NOB that may play a critical role in nitrogen removal in engineered and natural environments (Lücker *et al*., 2015). Recent 16S rRNA-based molecular surveys have identified *Nitrotoga*-like sequences in a surprisingly wide range of habitats, including: glacial soils (Pradhan *et al*., 2010; Sattin *et al*., 2009; Schmidt *et al*., 2009; Srinivas *et al*., 2011), an underground cave (Chen *et al*., 2009), a freshwater seep (Roden *et al*., 2012), drinking water (Kinnunen *et al*., 2017; White, DeBry, & Lytle, 2012), a subglacial Antarctic lake (Christner *et al*., 2014), Yellow Sea seawater (Na *et al*., 2011), salt marsh sediments (Martiny *et al*., 2011), rivers (Fan *et al*., 2016; Li *et al*., 2011), and various wastewater treatment plants (WWTPs) or sequence batch reactors (Bereschenko *et al*., 2010; Figdore, Stensel, & Winkler, 2017; Karkman *et al*., 2011; Lücker *et al*., 2015; Saunders *et al*., 2015; Yang, Chen, & Li, 2016; Ziegler *et al*., 2016). The relative abundance of *Nitrotoga*-like sequences in several WWTPs and a subglacial lake were as high as 2-13% of the total bacterial community, and were occasionally the only observed NOB (Christner *et al*., 2014; Lücker *et al*., 2015; Saunders *et al*., 2015). The distribution and relative abundance of *Nitrotoga*-like sequences suggest these organisms likely play a critical nitrogen cycling role in diverse environments; however, few studies have attempted to characterize their diversity or physiology.

Only four *Nitrotoga* cultures have been reported to date, with no confirmed isolates and no genome sequences are available within the genus. *Candidatus* Nitrotoga arctica was enriched from permafrost (Alawi *et al*., 2007); *Candidatus* Nitrotoga sp. HAM-1 from activated sludge (Alawi *et al*., 2009); *Candidatus* Nitrotoga sp. HW29 from an aquaculture system (Hüpeden *et al*., 2016); and *Candidatus* Nitrotoga sp. AM1 from coastal sand (Ishii *et al*., 2017). All *Nitrotoga* were enriched at low temperatures (4-17°C), however temperature optima were slightly higher (13-22°C) (Alawi *et al.*, 2007, 2009; Hüpeden *et al.*, 2016; Ishii *et al.*, 2017). *Ca.* Nitrotoga arctica has an intermediate nitrite affinity and is adapted to low nitrite concentrations (0.3 mM) compared to other NOB (Nowka *et al.*, 2015; Kits *et al.*, 2017).

Here, we describe near-complete draft genome sequences of four novel *Nitrotoga* species enriched from river water column and sediment samples. Each organism contained genomic capabilities for diverse metabolisms, which could support their growth in a wide range of habitats. This study represents the first reported cultivation of *Nitrotoga* from natural, freshwater systems and the first genome profiling from within the genus. These findings extend our understanding of freshwater nitrite oxidation and form the basis for further experimental work aimed at testing genomic predictions in culture and in the environment.

## METHODS

### Culture inoculation and growth

Two surface sediment samples were collected from the urban-impacted Cherry Creek in downtown Denver, CO (samples MKT and LAW). Two water column samples were collected from two agriculturally-impacted rivers near Greeley, CO (about 100 km North of Denver, CO) (samples CP45 from the Cache la Poudre River and SPKER from the South Platte River). Sediment and water column samples were inoculated into Freshwater Nitrite Oxidizer Medium (FNOM) with 0.3 mM nitrite and incubated at room temperature in the dark (Supplemental Note). Enrichment cultures were transferred to new media approximately every two weeks. Enrichment was enhanced by serial dilution, rapid transfers at the beginning of nitrite consumption, and low volume transfers (as low as 0.1% inoculum vol./vol.). Nitrite consumption was regularly monitored in the cultures using a Griess nitrite color reagent (Griess-Romijn van Eck, 1966). Nitrite oxidation rates were determined in triplicate for each enrichment culture (Supplemental Note).

### DNA extraction and sequencing

DNA was extracted from each culture at mid- to late-phase of exponential nitrite oxidation (Supplemental Note). Extracted DNA was sheared using a Covaris S220 (Covaris, Woburn, MA) and libraries were prepped with an insert size of 400 bp using an Ovation Ultralow System V2 (No. 0344) kit (Nugen, San Carlos, CA) by the University of Colorado Anschutz Medical Campus Genomics Core. DNA was sequenced on an Illumina HiSeq 2500 using V4 chemistry (Illumina, San Diego, CA) with 125 bp paired end reads.

### Metagenome and *Nitrotoga* genome assembly and annotation

Metagenomes from each enrichment culture were assembled with quality filtered and trimmed reads using MEGAHIT (Li *et al.*, 2014) and contigs were binned using MetaBAT (Kang *et al.*, 2015). *Nitrotoga* genomes were assembled iteratively with SPAdes v3.9.0 (Bankevich *et al.*, 2012) (Supplemental Note). Genome bin completeness and contamination estimates were calculated using the CheckM v1.0.11 lineage workflow (Parks *et al.*, 2015) (Supplemental Note).

Final assemblies were filtered to remove contigs <2 kb, all of which had very low and uneven coverage estimates. The *Nitrotoga* genomes were aligned and contigs reordered with progressiveMauve (Darling *et al.*, 2010), using the *Nitrotoga* species from the MKT culture as a reference due to its long contig length and simple assembly graph. Genomes were submitted to the DOE-JGI Microbial Genome Annotation Pipeline (MGAP) (Huntemann *et al.*, 2015) for final contig trimming of ambiguous and low-complexity sequences. Gene annotations were evaluated based on results from MGAP including COG, KEGG, Pfam, and TIGRfam assignments, InterPro Scan, IMG term assignments, and final protein product assignments (Huntemann *et al.*, 2015 and references within), as well as a BLASTP search of all predicted CDS against the nr database and annotation with KASS (Moriya *et al.*, 2007). RNAs were detected via MGAP with CRT, pilercr, tRNAscan, hmmsearch, BLASTN, and cmsearch (Huntemann *et al.*, 2015 and references within).

### *Nitrotoga* comparative genomics

The Anvi’o pangenome pipeline (Eren *et al.*, 2015) was used to cluster (mcl=10) coding sequences from each *Nitrotoga* genome in order to establish a ‘core genome’ of genes shared by all four *Nitrotoga* spp., and genes unique to each genome or shared by two or three *Nitrotoga* spp. Average nucleotide identity (ANI) and average amino acid identity (AAI) were calculated using the online enveomics tools (Rodriguez-R and Konstantinidis, 2016).

### *Nitrotoga* phylogenetic analyses

Near full-length *Nitrotoga*-like 16S rRNA gene sequences in the NCBI nt database were identified by a BLASTN search. Sequences were retained if they were ≥1300 bp in length and had ≥95% identity to any of the cultivated *Nitrotoga* 16S rRNA gene sequences from this study. 16S rRNA gene sequences from the BLASTN search were aligned with MAFFT (Katoh and Standley, 2013) and manually trimmed. A maximum likelihood tree was generated using RAxML (version 8.2.9) with 100 rapid bootstraps and the GTRGAMMA model of nucleotide substitution. A single monophyletic clade was extracted from the resulting phylogenetic tree, which included all known *Nitrotoga* 16S rRNA gene sequences. Neighboring clades included members of different genera. Selected outgroup sequences were added to the extracted sequences and then were realigned and trimmed, and a maximum likelihood tree was generated with 1000 rapid bootstraps.

Amino acid sequences (≥833 amino acids in length) of Type II DMSO reductase family enzymes were compiled based on previous research (Lücker *et al.*, 2010, 2013; Ngugi *et al.*, 2015) and a BLASTP search of the NCBI nr database. Sequences were aligned using MAFFT (Katoh and Standley, 2013) and manually trimmed. A maximum likelihood tree was built using RAxML with 1000 rapid bootstraps and the LG likelihood model of amino acid substitution (based on PROTGAMMAAUTO selection). All trees were visualized and annotated in iTOL (Letunic and Bork, 2016).

### Distribution of *Nitrotoga*-like 16S rRNA gene sequences in the environment

Full-length 16S rRNA gene sequences from each *Nitrotoga* culture were submitted to the IMNGS online server (Lagkouvardos *et al.*, 2016) for searches against all 183,153 16S rRNA gene amplicon runs from the NCBI Sequence Read Archive (SRA) (March 2018 release). A minimum identity threshold of 97% was chosen to represent *Nitrotoga*-like sequences at the genus level due to phylogenetic resolution of 16S rRNA gene sequences (see results). Samples were removed if they did not have at least 100 reads associated with *Nitrotoga*-like operational taxonomic units (OTUs). Data was summarized by SRA annotated environment categories. In some cases, SRA annotated categories were merged together: “aquatic”, “freshwater”, and “pond” were merged into “freshwater”; “freshwater_sediment” and “sediment” were merged into “sediment”; “biofilm” and “microbial_mat” were merged into “biofilm”; “soil”, “terrestrial”, and “peat” were combined into “soil”; and “plant”, “rhizosphere”, and “root” were merged into “plant-associated”.

### Relative abundance of *Nitrotoga* in creek sediments and water column

Water column and sediment samples were collected from 18 sites along Bear Creek, 18 sites along Cherry Creek, and four sites along the South Platte River at the confluences of Bear Creek and Cherry Creek in Denver, Colorado, USA in Fall 2016 (Supplemental Note). DNA extracts were sent to the University of Illinois Roy J. Carver Biotechnology Center, Urbana, Illinois for total 16S rRNA amplicon sequencing using the 515F-Y and 926R primers (Parada *et al.*, 2015; Quince *et al.*, 2011). Libraries were prepared with the Fluidigm 48.48 Access Array IFC platform (Fluidigm Corporation, South San Francisco, CA) as previously described (Ramanathan *et al.*, 2017), and sequenced on an Illumina HiSeq with Rapid 250 bp paired-end reads (Illumina, San Diego, CA). Filtered reads were clustered into OTUs at 97% sequence identity and taxonomy was assigned with a BLASTN search against the SILVA 16S rRNA gene database (release 128) (Quast *et al.*, 2013).

## RESULTS AND DISCUSSION

### Enrichment and nitrite oxidation

The four nitrite-oxidizing cultures were enriched for 17 (CP45 and SPKER) or 20 (LAW and MKT) months via serial dilution and rapid transfer. Nitrite oxidation rates (calculated across three measurements during logarithmic nitrite oxidation) averaged 135.6 ± 21.5 μM NO_2_^-^/day (Figure 1), which was similar to previously reported data for *Ca.* Nitrotoga arctica (Nowka *et al.*, 2015). 16S rRNA gene sequence analyses and PCR with NOB specific primers revealed that each culture contained only one NOB related to the *Candidatus* Nitrotoga genus (Betaproteobacteria class; Nitrosomonadales order; Gallionellaceae family).

**Figure 1.**
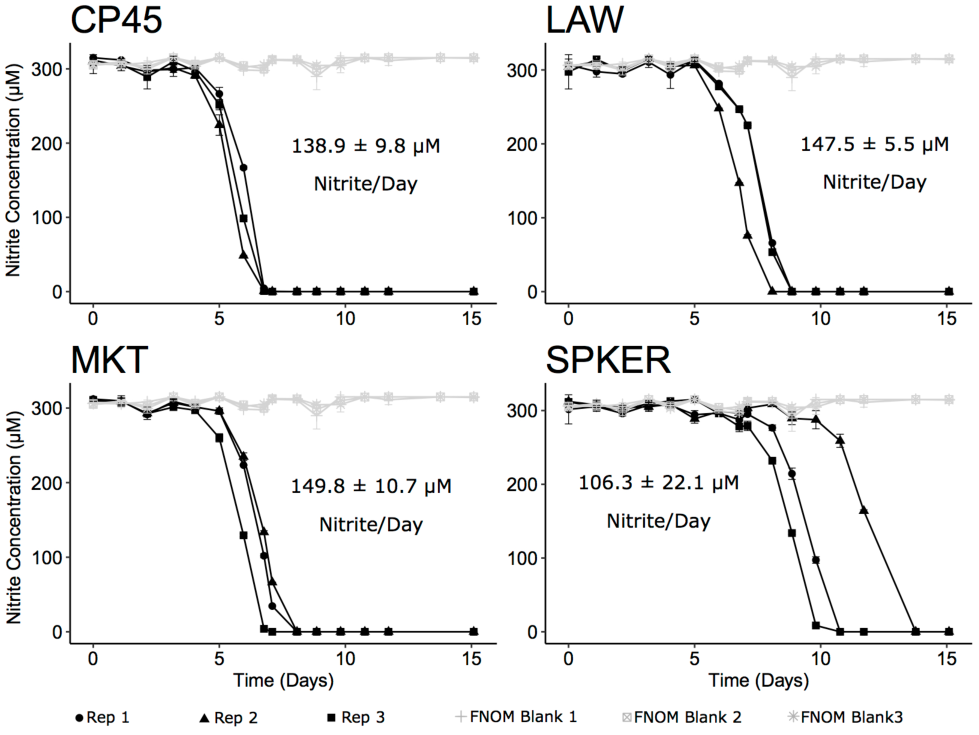
Nitrite consumption by *Nitrotoga* enrichment cultures over time. Each enrichment culture was inoculated in three replicates, and nitrite concentration was quantified colorimetrically in triplicate at each time point. Error bars show the standard deviation of each time point; error bars that appear to be missing are too small to be visualized. Sterile FNOM was used as a control and plotted with each culture. Logarithmic declines in nitrite concentration were used to calculate the nitrite oxidation rate for each biological replicate. The average nitrite consumption per day is shown with standard deviation among triplicate cultures.

We propose the following names for the enriched *Nitrotoga* species: *Candidatus* Nitrotoga mira MKT (n, nominative adjective, *mirus*, surprising) for the species enriched from Cherry Creek sediment and the unexpected enrichment of freshwater *Nitrotoga*; *Candidatus* Nitrotoga auraria LAW (f, nominative, noun, *aurarius*, of gold) for the species enriched from Cherry Creek sediment located near the historic Auraria settlement; *Candidatus* Nitrotoga amnis CP45 (m, genitive, noun, *amnis*, river) for the species enriched from Cache la Poudre River water; and *Candidatus* Nitrotoga coloradensis SPKER (f, adjective, Colorado, state central US) for the species enriched from water in the South Platte River, which flows through northeastern Colorado. Strain names represent sampling sites specific to each culture.

### Metagenome assembly and binning

Metagenomic sequencing of the enrichment cultures showed that each culture contained 8-32 genome bins (CP45: 8; MKT: 14; LAW: 23; SPKER: 32), likely corresponding to a similar number of species in each enrichment culture. Attempted co-assemblies of all four metagenomes were not successful as each enrichment culture had different species compositions. Each metagenome assembly contained only one predicted NOB related to *Nitrotoga* spp, along with other Proteobacteria including *Pseudomonas* spp, *Methylotenera* spp, and several members of the Comamonadaceae family (based on EMIRGE-assembled 16S rRNA genes (Miller *et al.*, 2011) and the CheckM lineage workflow). One genome bin from the SPKER metagenome likely belonged to a protozoan in the Neobodonida order that may prey on bacteria in culture.

### *Nitrotoga* genome assembly

The *Nitrotoga* genome bins from each culture had very high average coverage (201-398X). *Nitrotoga* genome bins were predicted to be 99.8% complete with ≤0.3% contamination by CheckM (after manual assignment of some marker genes; Supplemental Note; Supplementary Table S1). These high-quality draft sequences are the first known *Nitrotoga* genomes. The closest relatives with genome sequences were *Sideroxydans lithotrophicus* ES-1, *Gallionella capsiferriformans* ES-2, and *Ca.* Gallionella acididurans ShG14-8.

*Nitrotoga* genomes ranged in size from 2.707-2.982 Mbp, with 23-59 contigs and GC content between 47.5% and 48.8% (Supplementary Table S1). The number of coding sequences ranged from 2,574-2,858 with 36-39 tRNAs encoding all twenty amino acids. ANI values between the four *Nitrotoga* genomes ranged from 85.9%-93.4% (Table 1), while AAI values had a similar range of 87.5%-94.6%. These ANI values were indicative of each enrichment containing a novel *Nitrotoga* species, as they fell below the assumed 95% ANI threshold that separates most bacterial species (Caro-Quintero & Konstantinidis, 2012; Jain *et al*., 2017; Konstantinidis, Rosselló-móra, & Amann, 2017; Rodriguez-R & Konstantinidis, 2014). AAI values indicated these organisms were still highly conserved and likely shared many of the same traits. The SPKER *Nitrotoga* genome was considerably more divergent than the CP45, MKT, or LAW genomes.

**Table 1.** Average nucleotide identity (ANI) between enriched Nitrotoga genomes and close relative genomes available on NCBI (unhighlighted cells; top right) and average amino acid identity (AAI) pairwise comparisons (highlighted cells; bottom left).

|  | *Ca. Nitrotoga*<br>*amnis* CP45 | *Ca. Nitrotoga*<br>*auraria* LAW | *Ca. Nitrotoga*<br>*mira* MKT | *Ca. Nitrotoga*<br>*coloradensis*<br>SPKER | *Sideroxydans*<br>*lithotrophicus*<br>ES-1 | *Gallionella*<br>*capsiferriformans* ES-2 | *Ca. Gallionella*<br>*acididurans*<br>ShG14-8 | *Ferriphaseus*<br>*amnicola* | *Ferriphaseus*<br>sp. R-1 |
| --- | --- | --- | --- | --- | --- | --- | --- | --- | --- |
| *Ca. Nitrotoga*<br>*amnis* CP45 | 100.0% | 92.6% | 93.3% | 86.2% | 76.2% | 76.2% | 75.9% | 74.5% | 76.2% |
| *Ca. Nitrotoga*<br>*auraria* LAW | 93.8% | 100.0% | 93.4% | 86.2% | 74.2% | 74.3% | 74.8% | 73.8% | 74.1% |
| *Ca. Nitrotoga*<br>*mira* MKT | 93.9% | 94.6% | 100.0% | 85.9% | 75.6% | 76.5% | 76.9% | 75.7% | 76.6% |
| *Ca. Nitrotoga*<br>*coloradensis*<br>SPKER | 87.7% | 87.9% | 87.5% | 100.0% | 73.9% | 76.1% | 78.2% | 74.0% | 75.0% |
| *Sideroxydans*<br>*lithotrophicus*<br>ES-1 | 65.8% | 66.1% | 65.8% | 64.5% | 100.0% | 76.4% | 78.0% | 78.4% | 78.7% |
| *Gallionella*<br>*capsiferriformans* ES-2 | 63.2% | 64.2% | 63.7% | 63.3% | 66.1% | 100.0% | 76.7% | 75.9% | 78.2% |
| *Ca. Gallionella*<br>*acididurans*<br>ShG14-8 | 64.6% | 64.5% | 64.6% | 64.2% | 67.1% | 68.3% | 100.0% | 76.5% | 77.1% |
| *Ferriphaseus*<br>*amnicola* | 65.1% | 65.2% | 65.1% | 64.5% | 67.5% | 66.4% | 65.3% | 100.0% | 83.7% |
| *Ferriphaseus*<br>sp. R-1 | 65.0% | 65.3% | 65.6% | 64.6% | 67.9% | 66.0% | 65.5% | 84.7% | 100.0% |

Anvi’o pangenome analysis showed that coding sequences from all four genomes (10,666 in total) grouped into 4,001 protein clusters (PCs) (Figure 2). The core genome was represented by 1,803 PCs found in all four *Nitrotoga* genomes (45.1% of total). Each individual genome (e.g., MKT only) contained 293-625 unique PCs, while 566 PCs were shared among two or three genomes. Of the 2,198 accessory genome PCs, 1,041 were annotated as hypothetical proteins.

**Figure 2.**
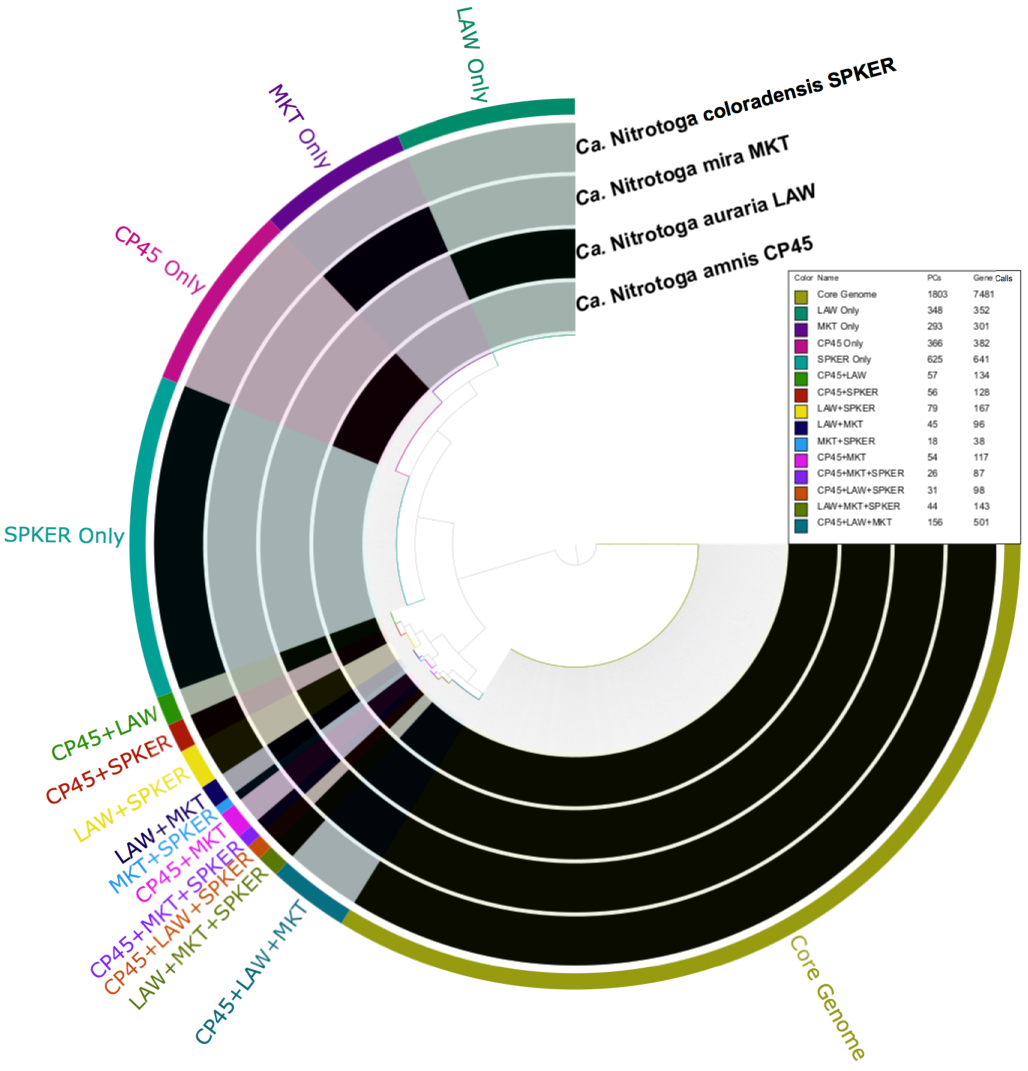
Anvi’o pangenome analysis of four *Nitrotoga* genomes. Coding sequences for all four genomes (10,666) in total grouped into 4,001 protein clusters (PCs), based on a pairwise BLAST of all coding sequences from all *Nitrotoga* genomes and clustering using the MCL algorithm (mcl=10). The core genome was represented by 1,803 PCs found in all four *Nitrotoga* genomes. Each individual genome (e.g., MKT only) contained 293-625 unique PCs not found in any other *Nitrotoga* genome.

Despite genomic level differences, the *Nitrotoga* 16S rRNA gene sequences were highly conserved (Figure 3; Supplementary Figure S1). *Nitrotoga* 16S rRNA genes were present at double coverage in all *Nitrotoga* genomes, indicating a gene duplication. The 16S rRNA gene from the SPKER genome had three single nucleotide variants (SNVs) present in ~50% of mapped reads, likely indicating small differences between the duplicate copies. A consensus sequence was used here, as the variants could not be isolated with paired reads. Among the four *Nitrotoga* 16S rRNA gene sequences assembled in this study, the most divergent are 99.4% identical across a 1,544 bp alignment (9 total mismatches). Pairwise comparisons of all 8 enriched *Nitrotoga* 16S rRNA gene sequences (4 previous studies, 4 from this study) average 99.5% identical across near full-length genes (Figure 3b) and a comparison of all 60 *Nitrotoga-* like sequences in Figure 3, comprising the known *Nitrotoga* genus, averaged 98.7% (Supplemental Figure 1). The highly conserved nature of *Nitrotoga* 16S rRNA gene sequences does not likely represent genome-level conservation across the genus, based on 16S rRNA:genome comparisons in the present study.

**Figure 3.**
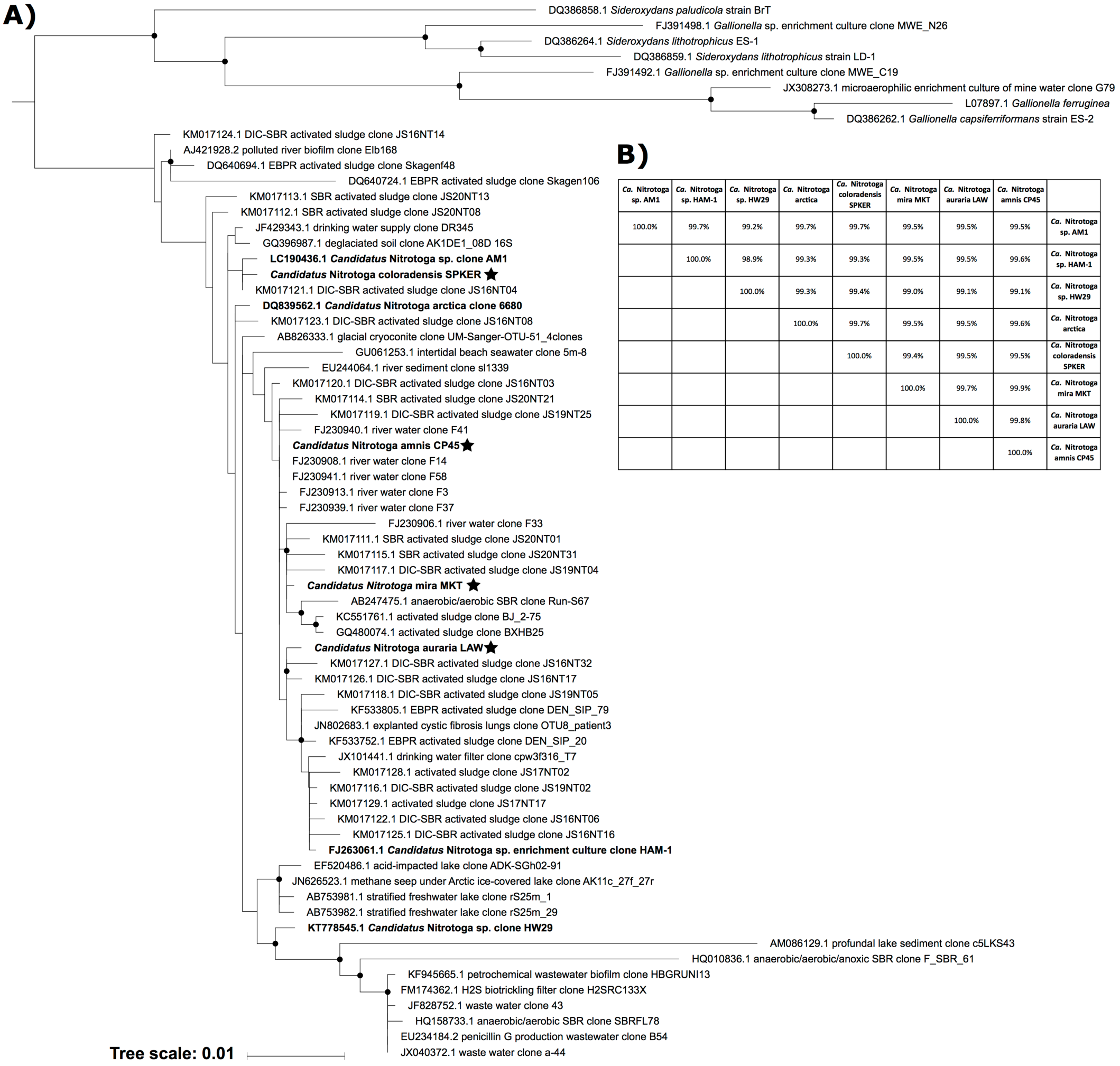
**A)** Maximum likelihood phylogenetic tree of 16S rRNA gene sequences from representative *Ca*. Nitrotoga sequences and close relatives. Sequences were aligned across 1,422 positions; phylogenetic trees were generated using RAxML (version 8.2.9) with 1,000 rapid bootstraps and the GTRGAMMA model of nucleotide substitution. Nodes with bootstrap support values ≥50% are shown with a black circle. A midpoint root was used when selecting outgroup sequences. Bolded sequence names have been enriched in culture. Nodes with a star represent organisms presented in this study. **B)** BLASTN comparisons of the near full-length 16S rRNA gene sequences from the eight known enriched *Nitrotoga* sp.

### *Nitrotoga* nitrogen metabolism

#### Nitrite oxidation

Nitrite oxidoreductase (NXR) is a heterotrimeric enzyme (NxrABC) that oxidizes nitrite to nitrate, liberating two electrons from a water molecule (Supplemental Note). NXR is a member of the Type II DMSO reductase family of molybdenum enzymes. NXR is bound to the cell membrane and can be classified into at least two distinct phylogenetic and functional groups based on orientation towards the cytoplasm or periplasm (Lücker *et al.*, 2010). Oxidation kinetics studies have associated NXR orientation with ecological niche formation, as bacteria with cytoplasmic-facing NXR (*Nitrobacter*, *Nitrococcus*, and *Nitrolancea*) typically dominate in relatively high nitrite environments over bacteria with periplasmic-facing NXR (*Nitrospira*, *Nitrospina*, and *Candidatus* Nitromaritima) (Kits *et al*., 2017; Koch *et al*., 2015; Lücker *et al*., 2010; Lücker, *et al*. 2013; Nowka *et al*., 2015; Sorokin *et al*., 2012; Spieck, *et al*. 1996; Spieck, *et al*. 1998; Starkenburg *et al*., 2006). Periplasmic-facing NXR have the energetic benefit of H^+^ release contributing to the proton motive force, while nitrite must be pumped into the cell for cytoplasmic-facing NXR.

*Nitrotoga nxr* genes were found on single contigs (6.7-9.5kb) within each genome forming an *nxrABC* operon, with an additional *nxrD* chaperone (Supplemental Note). Based on the number of neighboring contigs in the assembly and read coverage estimates, the *nxr* genes in the CP45, LAW, and MKT genomes are thought to be duplicated. The SPKER *nxr* operon is likely present three times throughout the genome based on the same criteria.

*Nitrotoga* NxrA amino acid sequences were divergent from all other members of the Type II DMSO reductase enzyme family, and the closest relatives were putative archaeal nitrate reductase (NarG) proteins (Figure 4a). The deeply branching *Nitrotoga* NxrA amino acid sequences may represent a fourth evolutionary development of nitrite oxidation, separate from the cytoplasmic-facing, periplasmic-facing, and phototrophic *Thiocapsa* nitrite oxidizers (Hemp *et al.*, 2016; Lücker *et al.*, 2010). Interestingly, some contigs surrounding the *Nitrotoga nxr* operons contained putative transposase and integrase genes, possibly suggesting that the *Nitrotoga nxr* genes were horizontally transferred similar to the theory of periplasmic-facing *nxr* gene development (Lücker *et al.*, 2010). The *Nitrotoga* genus also represents the only known Betaproteobacterial lineage of nitrite-oxidation.

**Figure 4.**
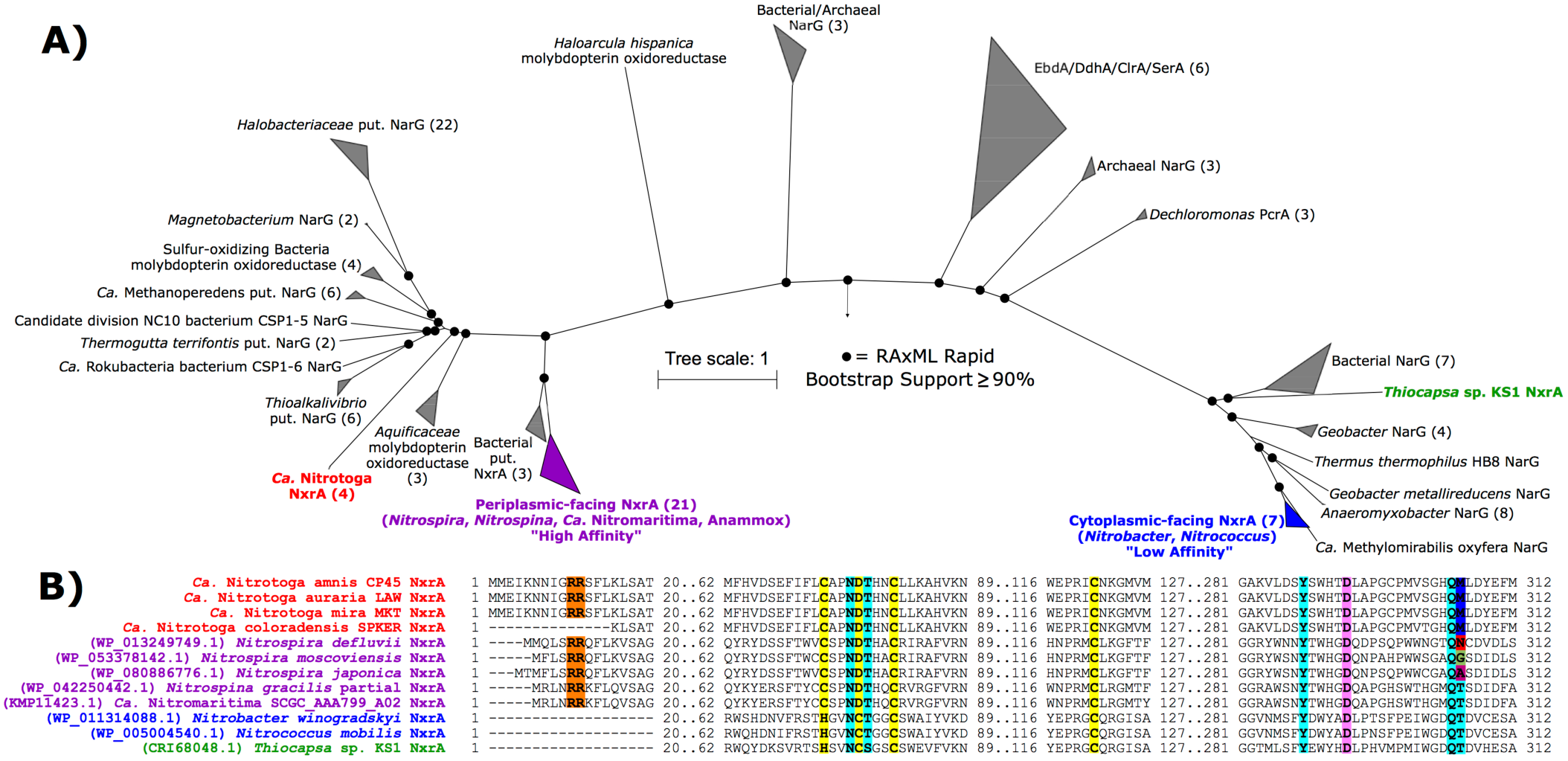
Phylogenetic and structural analysis of the alpha subunit of nitrite oxidoreductase (NxrA). **A)** Phylogeny of 122 members of the Type II DMSO reductase protein family aligned and manually trimmed to 1,651 amino acid positions. References were selected to include both cytoplasmic-facing “Low Affinity” and periplasmic-facing “High Affinity” NxrA (Ngugi *et al.*, 2015), as well as other family members: PcrA (perchlorate reductase), EbdA (ethylbenzene dehydrogenase), DdhA (dimethylsulfide dehydrogenase), ClrA (chlorate reductase), SerA (selenite reductase), and NarG (nitrate reductase). Putative enzymes are marked as “put.”. The number of sequences in collapsed nodes are shown in parentheses. RAxML rapid bootstrap support values ≥90% are marked with a black circle. **B)** Partial alignment of selected important residues in *Nitrotoga*, periplasmic-facing, and cytoplasmic-facing NxrA (including *Thiocapsa*). Highlights represent Tat signal peptides (Orange), Fe-S cluster binding residues (Yellow), molybdenum coordinating residues (Pink), and nitrite/nitrate binding residues (Light Blue).

*Nitrotoga* NxrA from the CP45, LAW, and MKT genomes had a predicted twin-arginine translocation (Tat) signal peptide on the N-terminus (Figure 4b), which supports the excretion of this enzyme into the periplasm (Sargent, 2007). The SPKER genome NxrA was lacking a signal peptide, although an alignment showed that the first 14 amino acids may be missing, as the entire peptide was otherwise aligned without gaps. The gene was located at the end of a contig and was missing a start codon.

Conserved residues were previously predicted to play a role in binding a molybdopterin cofactor, a [4Fe-4S] cluster, and the nitrite/nitrate substrate in NxrA and nitrate reductase NarG (Martinez-espinosa *et al.*, 2007; Lücker *et al.*, 2010). All *Nitrotoga* NxrA shared the conserved residues for the molybdopterin, iron-sulfur cluster, and four of the five nitrite/nitrate binding residues. However, an analysis of the fifth residue revealed a surprising lack of conservation among all NOB with at least five different residues predicted in known NxrA sequences, including variations within *nxrA* copies from a single genome (e.g., *Nitrospira nitrosa* has two *nxrA* copies, one with asparagine and one with glycine in this same position) (Figure 4b). This residue may not play a critical role in nitrite/nitrate binding, or may play a role in the variable nitrite oxidation kinetics observed previously (Nowka *et al.*, 2015; Kits *et al.*, 2017).

*Nitrotoga* NxrB were lacking a signal peptide, but may be excreted into the periplasm via a ‘hitch-hiker’ method as described previously (Lücker *et al.*, 2010; Martinez-espinosa *et al.*, 2007). All NxrB had coordinating residues for three [4Fe-4S] and one [3Fe-4S] cluster for electron conductance (Supplemental Figure 2).

One *Nitrotoga* NxrC predicted protein sequence was found in each genome with an N-terminal signal peptide for excretion. Signal peptides were previously observed in *Nitrospina gracilis* and *Candidatus* Nitromaritima NxrC (Lücker, *et al*. 2013; Ngugi, *et al*. 2015), however the predicted signal peptide cleavage site was located in the middle of a predicted transmembrane region. Biochemical studies confirmed the membrane-association of the NXR holoenzyme in *Nitrospina* and *Nitrospira* (Bartosch *et al*., 1999), indicating that NxrC were not fully translocated into the periplasm (Lücker *et al.*, 2010). *Nitrotoga* NxrC do not have predicted transmembrane regions, and are more similarly related to the soluble periplasmic gamma subunit of ethylbenzene dehydrogenase, which likely interact with cytochrome *c* proteins to shuttle electrons to the membrane (Kloer, *et al*. 2006); similar observations were made in the analysis of *Nitrospina gracilis* and *Candidatus* Nitromaritima genomes (Lücker, *et al*. 2013; Ngugi, *et al*. 2015).

Overall, sequence and phylogenetic analyses indicated that *Nitrotoga* possess a form of NXR that is divergent from known NOB. Phylogenetically, the alpha subunit was deeply branched near putative bacterial and archaeal NarG, but maintains all necessary residues for nitrite oxidation. The NxrA and NxrC subunits had signal peptides for excretion to the periplasm, but all subunits were lacking transmembrane domains for anchoring in the cytoplasmic membrane. This may suggest that *Nitrotoga* have a soluble NXR periplasmic holoenzyme, which has never been observed in NOB.

#### Dissimilatory and assimilatory nitrogen metabolism

All *Nitrotoga* genomes had genes for a NirK dissimilatory nitrite reductase for the reduction of nitrite to nitric oxide. *nirK* genes have been found in all other NOB genomes except *Nitrolancea hollandica* (Lücker *et al.*, 2013; Sorokin *et al.*, 2012), but their ultimate role is still unclear. CP45, LAW, and MKT genomes encoded a nitric oxide dioxygenase (*hmp*), which catalyzes the conversion of nitric oxide to nitrate, and is evolutionarily related to an O_2_-binding protein similar to hemoglobin (Gardner *et al.*, 1998). The role of *nirK* and *hmp* in *Nitrotoga* is unknown.

The *Nitrotoga* genomes encoded genes for transport of nitrite/nitrate (*narK*), formate/nitrite, and ammonium (*amtB*). The predicted periplasmic orientation of *Nitrotoga* NXR excludes the need for nitrite import into the cytosol, however genes for an assimilatory NirBD nitrite reductase and cytochrome c551/c552 to catalyze the reduction of nitrite to ammonia were found. *Nitrotoga* may also assimilate ammonia released during cyanide detoxification (Supplemental Note). A *Nitrotoga* enrichment culture from coastal sands grew faster when ammonium was added to the culture medium (Ishii *et al.*, 2017), likely due to the reduced need for assimilatory nitrite reduction.

### *Nitrotoga* energy metabolism and reverse electron flow

#### Nitrogen energetics

A complete electron transport chain was present in the *Nitrotoga* genomes (Supplemental Figure 3). After electrons are passed from NXR to cytochrome *c*, they are transferred to oxygen via a terminal oxidase (Complex IV). All *Nitrotoga* genomes contained genes for a *cbb_3_*-type cytochrome *c* oxidase, a member of the C-class heme-copper oxidases with an exceptionally high affinity for oxygen (Morris and Schmidt, 2013). These genes did not form a distinct operon, however there were no other candidate terminal oxidase genes in most *Nitrotoga* genomes (see below). Organisms possessing *cbb_3_*-type oxidases, including the NOB *Nitrospina gracilis* and the phototrophic nitrite-oxidizer *Thiocapsa* KS1, are likely capable of growth in microoxic environments (Han *et al.*, 2011; Lücker *et al.*, 2013; Hemp *et al.*, 2016). For instance, *Nitrospina*-like bacteria have been found to play a crucial role in carbon fixation in marine oxygen minimum zones, due in part to their *cbb3*-type terminal oxidases (Füssel *et al.*, 2012, 2017;Pachiadaki *et al.*, 2017). The possession of a *cbb_3_*-type terminal oxidase indicates that *Nitrotoga* species likely continue aerobic metabolisms at nanomolar O_2_ concentrations, allowing an incredibly wide habitat range (e.g., sediments, biofilms, marshes). *Nitrotoga* may also use alternative metabolisms (e.g., sulfur oxidation; see below) in low oxygen environments similar to *Nitrococcus mobilis* (Füssel *et al.*, 2017). The *Nitrotoga* genomes had several O_2_ binding proteins including protoglobin, hemerythrin, and potentially nitric oxide dioxygenase, which may facilitate survival under low oxygen conditions.

Intriguingly, the SPKER genome contained an additional *bd*-type terminal cytochrome *c* oxidase, which was also observed in the genomes of two close relatives: *Sideroxydans lithotrophicus* ES-1 and *Gallionella capsiferriformans* ES-2 (Emerson *et al.*, 2013). This terminal oxidase may serve an additional role in energy conservation from organic carbon sources (see below), as *bd*-type terminal oxidases only receive electrons from the quinone pool, while *cbb_3_*-type terminal oxidases can receive electrons from cytochromes or the quinone pool (Borisov *et al.*, 2011; Morris and Schmidt, 2013).

An F-type ATPase (Complex V) was present in all four *Nitrotoga* genomes for ATP generation via the proton motive force. The periplasmic oxidation of nitrite contributes to the proton gradient, as two protons are released in the reaction. Inorganic phosphate for ATP generation can be stored as polyphosphate in *Nitrotoga* and released by polyphosphate kinase and inorganic pyrophosphatase enzymes.

When nitrite is the only energy source, *Nitrotoga* must generate NADH via reverse electron flow to support normal cellular processes. A canonical cytochrome *bc_1_* (Complex III), typically found in proteobacteria, was missing from *Nitrotoga* genomes. However, a suite of genes encoding an alternative complex III (*actAB1B2CDEF*) (Refojo *et al.*, 2012) was found in all *Nitrotoga* genomes and in the related iron-oxidizing bacteria of the Gallionellaceae family (Emerson *et al.*, 2013). Electrons are distributed from the alternative complex III via quinones to succinate dehydrogenase (Complex II) for biochemical intermediate production, or to NADH dehydrogenase (Complex I) for the reduction of NAD^+^ (Supplemental Figure 3).

In addition to nitrite oxidation, NXR mediated nitrate reduction with electrons derived from alternative donors has been observed in members of the *Nitrobacter* and *Nitrospira* when oxygen is excluded (Bock *et al*., 1990; Koch *et al*., 2015; Sundermeyer-Klinger *et al*., 1984). Given the genomic potential for survival in low-oxygen environments, we predict that *Nitrotoga* are also capable of nitrate reduction via NXR when oxygen is not available. Alternative electron donors including organic carbon, reduced sulfur compounds, or hydrogen gas (see below) could contribute to the electron transport with NXR functionally replacing the terminal oxidase (Supplemental Figure 3).

#### Sulfur energetics

Genomic pathways indicated that *Nitrotoga* may be capable of sulfur oxidation. The recently described *Nitrococcus mobilis* Nb-231 genome encoded genes for aerobic sulfur oxidation, and their activity was confirmed in pure culture (Füssel *et al.*, 2017). *Nitrotoga* could likely harness the same sulfur sources using a periplasmic sulfite dehydrogenase (SOR) to oxidize sulfite (SO_3_^-^) to sulfate (SO_4_^2-^) with electron donation to cytochrome *c*, and a sulfide:quinone oxidoreductase (SQR) that couples hydrogen sulfide (H_2_S) oxidation to elemental sulfur (S^0^) and reduction of quinone (Supplemental Figure 3).

#### Hydrogen energetics

The catalytic subunit of a Group 3d [NiFe]-hydrogenase was identified in the CP45, LAW, and MKT genomes with the HydDB online tool (Søndergaard *et al.*, 2016). All other requisite genes were found nearby in the respective genomes. Group 3d [NiFe]-hydrogenases typically act as an NADH oxidoreductase, reducing NAD^+^ to NADH with concomitant oxidation of H_2_ to water (Peters *et al.*, 2015). Many sequenced NOB have a Group 3b [NiFe]-hydrogenase (Daims *et al*., 2015; Füssel *et al*., 2017; Lücker *et al*., 2013) that is also capable of elemental sulfur or polysulfide reduction to H_2_S, an advantageous enzyme that is lacking in *Nitrotoga* genomes. *Nitrospira moscoviensis* utilizes a Group 2a [NiFe]-hydrogenase, and growth on H_2_ was confirmed in culture (Koch *et al.*, 2016). Hydrogen oxidation could serve as yet another energy metabolism for *Nitrotoga* cells.

### *Nitrotoga* carbon metabolism

#### Carbon fixation

Nitrotoga genomes had genes for the complete Calvin cycle (Supplemental Note), supporting CO_2_ fixation for autotrophic growth. A sedoheptulose-bisphosphatase gene, an important intermediate enzyme needed to regenerate ribulose-1,5-bisphosphate, was missing but is also missing in the NOB *Nitrobacter winogradskyi*, and the ammonia-oxidizing betaproteobacteria *Nitrosomonas europaea*, and *Nitrosospira multiformis* (Chain *et al.*, 2003; Norton *et al.*, 2008; Starkenburg *et al.*, 2006). Fructose 1,6-bisphosphatase, which typically plays a role in gluconeogenesis, is thought to fill the same role in organisms lacking sedoheptulose-bisphosphatase (Wei *et al.*, 2004; Yoo and Bowien, 1995) and was present in *Nitrotoga* genomes. *Nitrolancea*, *Nitrococcus*, and *Nitrobacter* also utilize the Calvin cycle (Sorokin *et al.*, 2012; Füssel *et al.*, 2017; Starkenburg *et al.*, 2008).

#### Organic carbon utilization

*Nitrotoga* genomes encoded all genes for glycolysis, gluconeogenesis, the pentose phosphate pathway, and the tricarboxylic acid (TCA) cycle (Figure 5a). All necessary biochemical intermediates could be generated via these pathways, whether carbon is fixed via the Calvin cycle or imported into the cell. *Nitrotoga* genomes had genes for polysaccharide storage and exopolysaccharide synthesis, which may facilitate growth in biofilms (Supplemental Note).

**Figure 5.**
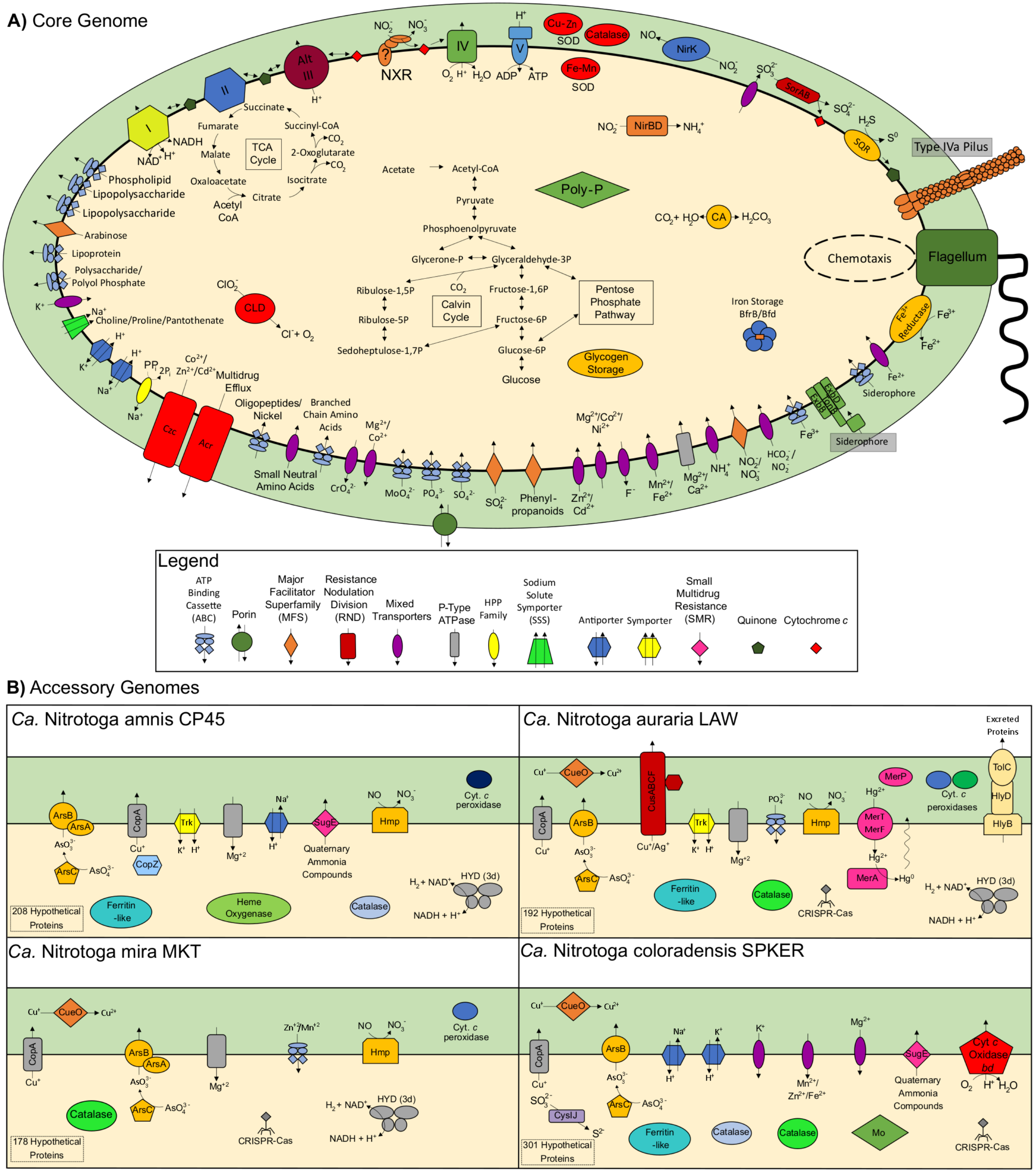
Schematic representation of *Nitrotoga* genome features. **A)** The core *Nitrotoga* genome including predicted functions shared by all four genomes. The question mark located in the NxrC subunit symbolizes the uncertainty in whether or not the holoenzyme is anchored to the cell membrane. **B)** Predicted function of individual genomic features found within one or more genomes, but not shared among all four. Each genome had 178-301 unique hypothetical proteins, and an additional 171 hypothetical proteins were shared among two or three genomes.

Various sugar transporters are present in many NOB, including *Nitrococcus mobilis*, *Nitrospira inopinata*, *Nitrospina gracilis*, and *Nitrolancea hollandica* (Füssel *et al.*, 2017; Daims *et al.*, 2015; Lücker *et al.*, 2013; Sorokin *et al.*, 2012), but none were identified in *Nitrotoga* genomes. However, several genes involved in the phosphotransferase system (PTS) to import and phosphorylate sugars were found. The PTS was also found in *Nitrobacter winogradskyi* and *Thiocapsa* sp. KS1 (Starkenburg *et al.*, 2006; Hemp *et al.*, 2016). EI and HPr subunits of the PTS were both found in *Nitrotoga*, but a complete set of substrate-specific EIIA, EIIB, and EIIC components were not identified. Only ascorbate-specific EIIA and nitrogen regulatory EIIA subunits were found. *Nitrotoga* genomes did possess relatives of a recently purified ABC transporter (AfuABC) that was found to import phosphorylated sugars instead of iron as originally reported (Sit *et al.*, 2015).

*Nitrotoga* genomes had genes for an acetyl-CoA synthetase, which produces acetyl-CoA from acetate, suggesting the use of small organic carbon molecules if they can enter the cytosol. The incomplete *Nitrotoga* genomes, as well as otherwise complete organic carbon oxidation pathways, leaves potential for future identification of organic carbon transport. Preliminary physiology tests showed a dramatically increased rate of nitrite oxidation when acetate and dextrose were provided to the cultures (data not shown).

### *Nitrotoga* iron acquisition

Iron is often a limiting nutrient in oligotrophic freshwater environments. *Nitrotoga* genomes encoded an impressive array of genes useful for iron scavenging, including a complete TonB-dependent transport system with up to four copies of the outer membrane transport energization protein complex (ExbB/ExbD/TonB) and up to 16 TonB-dependent outer membrane siderophore receptors (CP45: 10; LAW: 7; MKT: 6; SPKER: 16). All of the TonB-dependent transporters (TBDTs) fell within the CirA and/or Fiu superfamilies (based on BLASTP and conserved domain results), which have been shown to transport a wide variety of siderophores including citrate, aerobactin, enterobactin, and salmochelin (Porcheron et al., 2013). Eight TBDTs (MKT: 0; CP45: 1; LAW: 1; SPKER: 5) were preceded by FecI-FecR transcription factors known to monitor ferric dicitrate concentrations within the cell and control transcription of TBDTs in *E. coli* (Braun *et al.*, 2003). Siderophore synthesis seems unlikely within *Nitrotoga* (based on gene predictions; Supplemental Note), however *Nitrotoga* may scavenge siderophores from other bacteria within the community. Additionally, several iron pumps and iron storage proteins were present (Supplemental Note). Iron is a critical cofactor in the NXR enzyme and the varied iron acquisition methods of *Nitrotoga* may offer a competitive advantage for these organisms in iron-limited environments.

### *Nitrotoga* heavy metal transport and defense

Molybdate (MoO_4_^2-^) transport genes (*modABC*) were found in the CP45, LAW, and MKT genomes; while the SPKER genome had two *modA* copies but no *modB* or *modC*. However, the SPKER genome encoded a pair of molybdenum storage genes (*mosAB*) for intracellular storage of molybdenum (Fenske *et al.*, 2005), not found in other *Nitrotoga*. Molybdate is the naturally occurring form of molybdenum, which is a necessary cofactor for NXR, so *Nitrotoga* cells must have at least one method of acquiring molybdenum.

Heavy metal efflux systems were prevalent in all *Nitrotoga* genomes. An *apaG* cobalt and magnesium efflux protein, as well as complete cobalt-zinc-cadmium resistance transporters (*czcABC*) were found in all genomes. A chromate transporter and divalent cation tolerance protein may help protect against heavy metal accumulation in *Nitrotoga*.

Arsenate (AsO_4_^3-^) can enter cells through normal phosphate transport systems, and *Nitrotoga* had two different mechanisms for arsenate removal (Figure 5b). The LAW and SPKER genomes had *arsC* and *arsB* genes responsible for reducing arsenate to arsenite (AsO_3_^3-^), and pumping arsenite from the cytoplasm, respectively. The CP45 and MKT genomes encoded an additional subunit, *arsA*, which acts as an ATPase to provide energy for export, while the ArsB pump can work alone with energy from the proton motive force.

All *Nitrotoga* genomes encoded the alpha (*cusA*) and beta (*cusB*) portions of the Cus Cu(I)/Ag(I) efflux system, composing the inner membrane transporter and periplasmic membrane fusion protein, respectively. The outer membrane factor channel protein, *cusC*, was only found in a complete operon in the LAW genome along with the *cusF* metallochaperone and *cusR*-*cusS* two-component signaling system. The MKT and SPKER genomes had *cusR* and *cusS* genes preceding a TolC-like outer membrane protein that could be CusC, but these were found separate from *cusAB*. A second copper exporter belonging to the Cue system was found in the LAW, MKT, and SPKER genomes consisting of a Cu^2+^ exporter (*copA*) and multicopper oxidase (*cueO*). The CP45 genome alone had a *copZ* gene that serves as a chaperone for Cu^+^ export via CopA.

Chromate (CrO_4_^2-^) can enter cells via normal sulfate uptake systems. A putative soluble chromate reductase was found in all *Nitrotoga* genomes that can reduce the Cr(VI) found in chromate to the less toxic Cr(III), producing harmful reactive oxygen species (ROS) in the process which must be dealt with by other defenses (Thatoi *et al*., 2014). In addition, a chromate transporter gene (*chrA*) was found in all genomes for the excretion of chromate ions from the cytoplasm.

A variety of other heavy metal defenses were found in individual *Nitrotoga* genomes (Figure 5b), for instance, the CP45 genome encoded a gold/copper resistance efflux pump belonging to the resistance nodulation division (RND) superfamily. The LAW genome encoded a mercury resistance system (*merRTPFA*) for Hg(II) transport and reduction to volatile Hg(0), and the LAW and SPKER genomes encoded *terC*-like proteins associated with tellurite resistance but appear to be lacking other required genes for tellurite efflux and/or reduction. Heavy metal transport is critical for cofactor acquisition (e.g., iron, molybdenum) and for defense against metals that are toxic at even low concentrations (e.g., mercury, arsenic). *Nitrotoga* genomes possessed varied metal transporters, which likely help support cells in the contaminated rivers sampled in this study.

### Other defense mechanisms in *Nitrotoga*

In support of the aerobic metabolisms of *Nitrotoga*, all four genomes contained necessary catalase, superoxide dismutase (Cu-Zn and Fe-Mn families), and peroxiredoxin genes to combat reactive oxygen species (ROS) (Figure 5). Cytochrome *c* peroxidase genes were found in the CP45, LAW, and MKT genomes to intercept peroxides in the periplasm with electrons donated from cytochrome *c*. Additionally, a rubrerythrin protein used to combat ROS specifically in anaerobic bacteria was found, a reminder of the likely microaerophilic or anaerobic ancestry of *Nitrotoga* within the Gallionellaceae.

A few antibiotic resistance genes were present among the *Nitrotoga* genomes, including a generic antibiotic biosynthesis monooxygenase, VanZ-like family proteins that may confer low-level antibiotic resistance, and a broad-spectrum multidrug efflux system (AcrAB-TolC). An erythromycin esterase homolog was found in the LAW genome, which may confer resistance to erythromycin. Preliminary physiology tests indicated that *Nitrotoga* continued nitrite oxidation in the presence of various antibiotics (data not shown).

The SPKER *Nitrotoga* genome possessed an interesting suite of genes responsible for DNA phosphorothionation (*iscS*, *dndBCDE*) (swapping a non-bridging oxygen atom for a sulfur atom) that offers resistance to a related restriction-modification system (*dptHGF*) (Xu *et al*., 2010) found nearby on the genome. DNA phosphorothionation was recently found to expand microbial growth range under multiple stresses due to its intrinsic antioxidant function (Yang *et al.*, 2017). This ability may greatly expand the environments in which *Ca.* Nitrotoga coloradensis SPKER can proliferate, adding to its already wide potential range.

### Motility and chemotaxis

All *Nitrotoga* genomes carried genes necessary for flagellar assembly and operation, as well as signal transducing pathways to stimulate gliding or twitching motility via a type IV pilus assembly. All genomes had general chemotaxis genes, but the SPKER genome was notably missing an aerotaxis receptor (*aer*) capable of detecting oxygen concentrations. If *Nitrotoga* are indeed motile, they may migrate towards areas of greater reducing potential, and likely lose external motile structures when growing in biofilms or in culture, as the presence of flagella or pili have not been noted in previous studies (Alawi *et al.*, 2007, 2009; Hüpeden *et al.*, 2016; Lücker *et al.*, 2015; Ishii *et al.*, 2017).

The presence of a single acyl homoserine lactone synthase (LuxI homolog) in each genome, as well as several LuxR family transcriptional receptors and multiple homoserine efflux transporters (RhtA), indicate that *Nitrotoga* may use quorum sensing to communicate. Other NOB have been documented to use quorum sensing at high cell densities and the *Nitrotoga* LuxI homolog grouped phylogenetically with other sequenced Betaproteobacteria (Sayavedra-Soto *et al.*, 2015; Mellbye *et al.*, 2017). *Nitrotoga* have been documented to grow in flocs and were shown to co-aggregate with AOB in WWTPs (Alawi *et al*., 2007; Lücker *et al*., 2015), so they may use quorum sensing to form microcolonies in response to their environment.

### Environmental distribution

*Nitrotoga*-like 16S rRNA sequences were present in 2,410 different samples (from 183,154 total SRA runs analyzed with IMNGS) (Figure 6). *Nitrotoga* containing samples comprised 70 different user-defined environments with a global distribution across seven continents spanning from the tropics to the poles (Figure 6C). About 9% of all freshwater environments (517/5,678) had *Nitrotoga*-like OTUs, with some communities consisting of nearly 10% relative abundance of *Nitrotoga*-like OTUs. Nearly half of all wetland samples (24/49) had *Nitrotoga*-like OTUs with an average relative abundance of 0.89%, however most of these samples were part of the same BioProject. A high proportion of activated sludge (15%) and wastewater (17%) samples also harbored relevant populations of *Nitrotoga*-like OTUs (0.55% average relative abundance), similar to previous reports (Lücker et al., 2015). The relative abundance of *Nitrotoga*-like OTUs ranged from 0.02%-13.03% across 540 soil samples. Of the few (6/6,233) marine samples that had *Nitrotoga*-like OTUs, most were found near freshwater rivers or wetlands.

**Figure 6.**
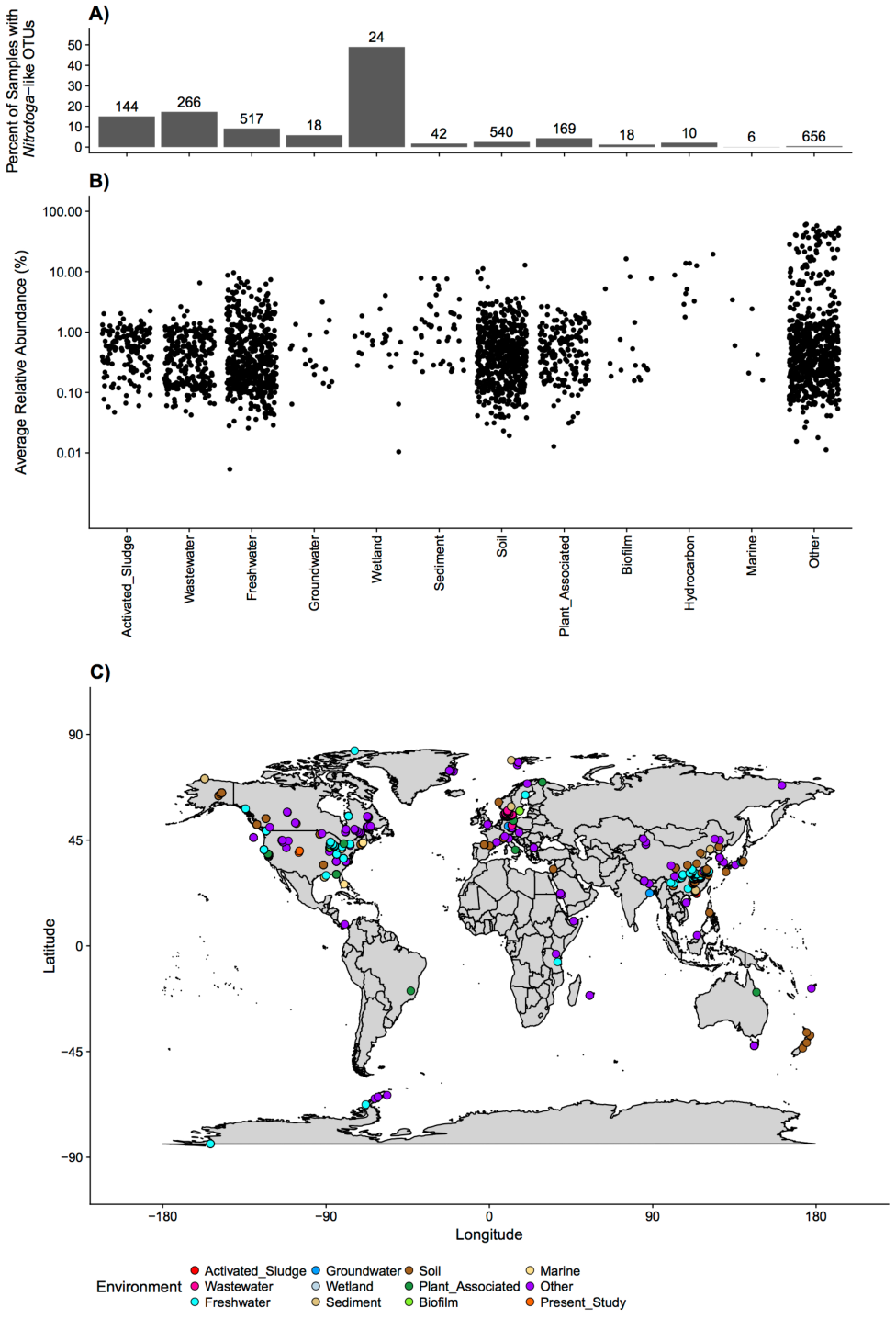
Global distribution of *Nitrotoga*-like sequences. OTUs from 16S rRNA gene amplicon studies deposited as SRA runs were clustered by IMNGS. All runs with OTUs ≥ 97% identity to cultured *Nitrotoga* 16S rRNA gene sequences (from this study), an alignment of at least 200 bp, and at least 100 total reads were kept. **A)** The percent of SRA runs with *Nitrotoga*-like OTUs within respective environments are shown with the number of SRA runs with *Nitrotoga*-like OTUs displayed above each bar. **B)** The relative abundance of *Nitrotoga*-like OTUs was averaged across all four queried *Nitrotoga* 16S rRNA gene sequences and plotted by environment. **C)** Global distribution of SRA runs with *Nitrotoga*-like OTUs. 484 of the 2,410 SRA runs with *Nitrotoga*-like OTUs did not have geographic information available, including all of the “Hydrocarbon” environmental samples. Orange points represent sampling locations for the enrichment cultures presented in this study.

A few samples in the SRA dataset had very high relative abundance of *Nitrotoga*-like OTUs compared to other samples. Several of these samples were associated with enrichment cultures from their respective environments (e.g., biofilm, hydrocarbon, and other). For example, a series of samples from a single study which involved enrichment from waterworks sand filters in the Netherlands had *Nitrotoga*-like OTUs comprising as much as 61% of the microbial community.

*Nitrotoga* distribution was also evaluated across 80 water column and sediment samples from three rivers in Colorado (Bear Creek, Cherry Creek, and South Platte River), based on amplification with general 16S rRNA gene primers targeting the total bacterial community (Supplemental Figure S4). *Nitrotoga*-like OTUs were identified in 85% of the samples, and *Nitrospira*-like OTUs were identified in 98% of the samples. Both groups co-occurred in 85% of the samples. No other NOB sequences were identified. Within each individual sample, the summed relative abundance ranged from 0-4.5% of the total bacterial community for *Nitrotoga*-like OTUs and 0-6.4% for *Nitrospira*-like OTUs. The summed relative abundance of *Nitrotoga*-like OTUs was greater than that of *Nitrospira*-like OTUs in 21% of the samples. *Nitrobacter* and *Nitrospira* are typically the more commonly studied NOB in freshwater systems (Daims *et al.*, 2016; Cai *et al.*, 2018), but future efforts should now also consider *Nitrotoga* as potential important players in freshwater nitrite oxidation.

The *Nitrotoga* species presented here were enriched from the water column and sediments of urban- and agriculturally-impacted rivers, suggesting a broad habitat range within freshwater systems and a greater role in freshwater nitrification than previously expected. Sampling for these *Nitrotoga* enrichment cultures occurred in winter and spring when *in situ* water temperatures are typically <15°C and cultures were grown with 300 μM nitrite. All other reported *Nitrotoga* were enriched at low temperatures (4-17°C) and low nitrite concentration (300 μM) (Alawi *et al.*, 2007, 2009; Hüpeden *et al.*, 2016; Ishii *et al.*, 2017). Temperature and substrate availability may play a role in niche differentiation of NOB, with *Nitrotoga* increasing in abundance at colder temperatures and lower nitrite concentrations, but further molecular and physiology experiments would be needed to confirm this.

## CONCLUSIONS

The enrichment of four novel *Nitrotoga* species, characterization of the first *Nitrotoga* genomes, and analysis of the distribution of *Nitrotoga* 16S rRNA sequences has expanded our knowledge of NOB ecology. The divergent NXR enzyme may indicate a novel evolution of nitrite oxidation in *Nitrotoga* separate from other known NOB. *Nitrotoga* have an exceptionally diverse metabolic profile, likely allowing proliferation under variable nutrient and oxygen conditions. The prevalence of *Nitrotoga*-like sequences found in globally distributed habitats (e.g., freshwater, soil, wetland, wastewater) suggests that *Nitrotoga* likely play a previously underappreciated functional role on a global scale. Future work should determine *Nitrotoga* ecophysiology, including nitrite oxidation kinetics and optimal growth conditions. Determining the functional redundancy and niche differentiation of *Nitrotoga* compared to other NOB will be important for understanding their role in nitrite oxidation under varying environmental conditions. Understanding the sensitivity or resilience of *Nitrotoga* to disturbances will be important for predicting their response to environmental change.

## ACKNOWLEGEMENTS

We thank Adrienne Narrowe and Christopher Miller for guidance on bioinformatic data analyses. We would also like to thank Hannah Clark, Nicklaus Deevers, Colin Beacom, Michael Kain, and Munira Lantz for their assistance in cultivation and preliminary physiology experiments and Sladjana Subotic and Anna Scopp for their help with environmental distribution sample collection and processing.

## COMPETING INTERESTS

The authors declare no competing interests. Portions of this manuscript were previously published as a part of University of Colorado Denver Master’s thesis submission (AB, 2017). Funding was provided by the University of Colorado, Denver and the City and County of Denver.

## SUPPLEMENTARY INFORMATION

Supplementary Information accompanies this paper.

---

**Supplemental Table 1.** Assembly statistics overview for *Nitrotoga* genomes. Percent completeness and contamination were estimated using CheckM. The asterisks indicate manual assignment of some marker genes used in completeness estimates (see Supplemental Note).

|  | **Num.**<br>**Contigs** | **GC %** | **N50 (bp)** | **Median Contig**<br>**Length (bp)** | **Total Length**<br>**(bp)** | **CDS** | **tRNA** | **rRNA** | **% Complete\*** | **%**<br>**Contamination** |
| --- | --- | --- | --- | --- | --- | --- | --- | --- | --- | --- |
| **Ca. *Nitrotoga***<br>**amnis CP45** | 59 | 48.8% | 69,731 | 35,034 | 2,816,237 | 2,665 | 36 | 6 | 99.8% | 0.30% |
| **Ca. *Nitrotoga***<br>**auraria LAW** | 33 | 48.7% | 123,222 | 58,971 | 2,814,534 | 2,730 | 37 | 6 | 99.8% | 0.30% |
| **Ca. *Nitrotoga***<br>**mira MKT** | 23 | 48.6% | 214,834 | 70,433 | 2,707,194 | 2,574 | 39 | 6 | 99.8% | 0.30% |
| **Ca. *Nitrotoga***<br>**coloradensis**<br>**SPKER** | 32 | 47.5% | 168,712 | 48,253 | 2,982,257 | 2,858 | 39 | 6 | 99.8% | 0.24% |

**Supplemental Figure S1.**
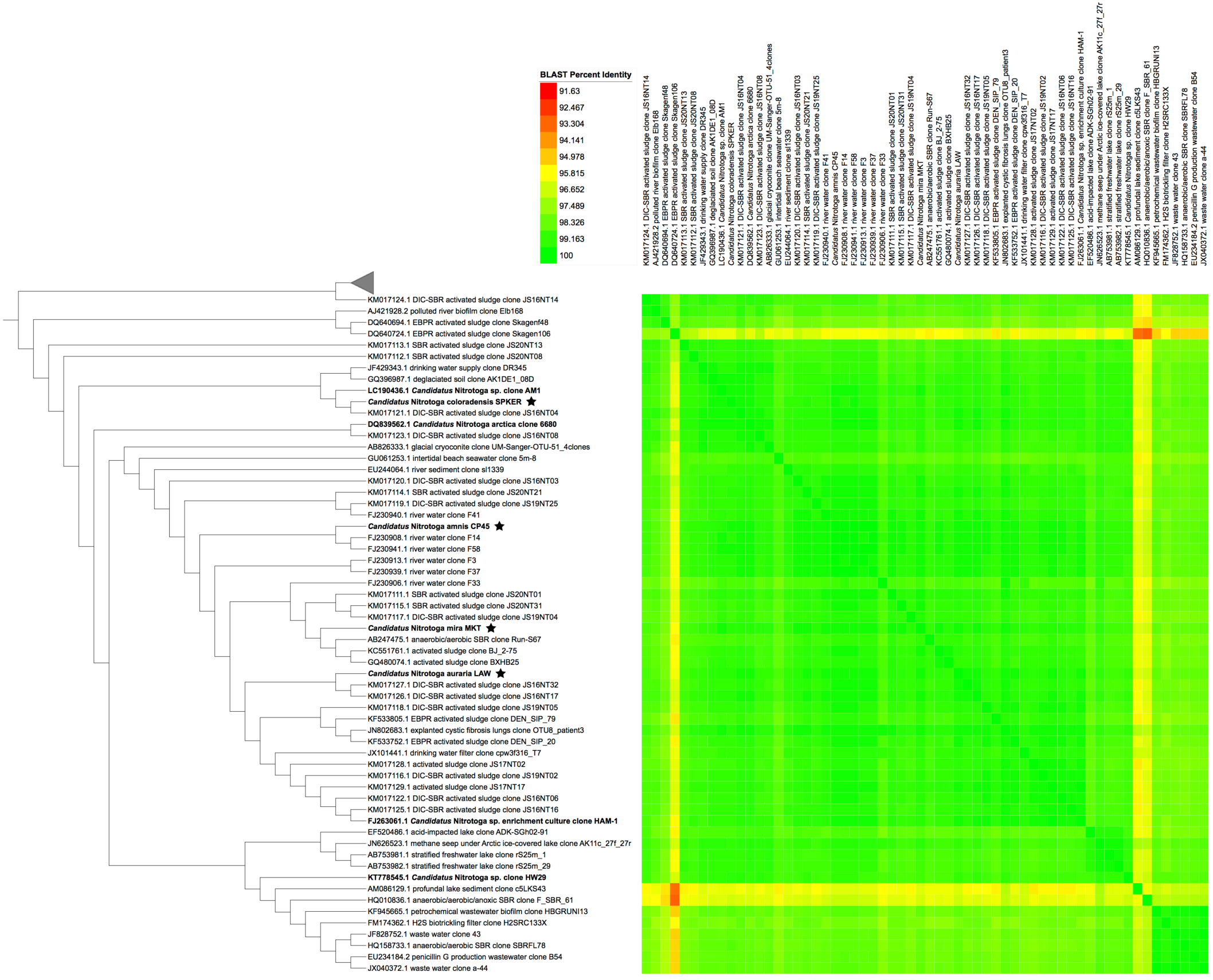
Pairwise BLAST comparisons of 60 *Nitrotoga* and *Nitrotoga*-like 16S rRNA gene sequences ≥1,300 bp presented as a heatmap. The phylogeny from Figure 3 is displayed on the left with branch lengths ignored. Bolded sequence names have been enriched in culture. Nodes with a star represent organisms presented in this study.

**Supplemental Figure S2.**
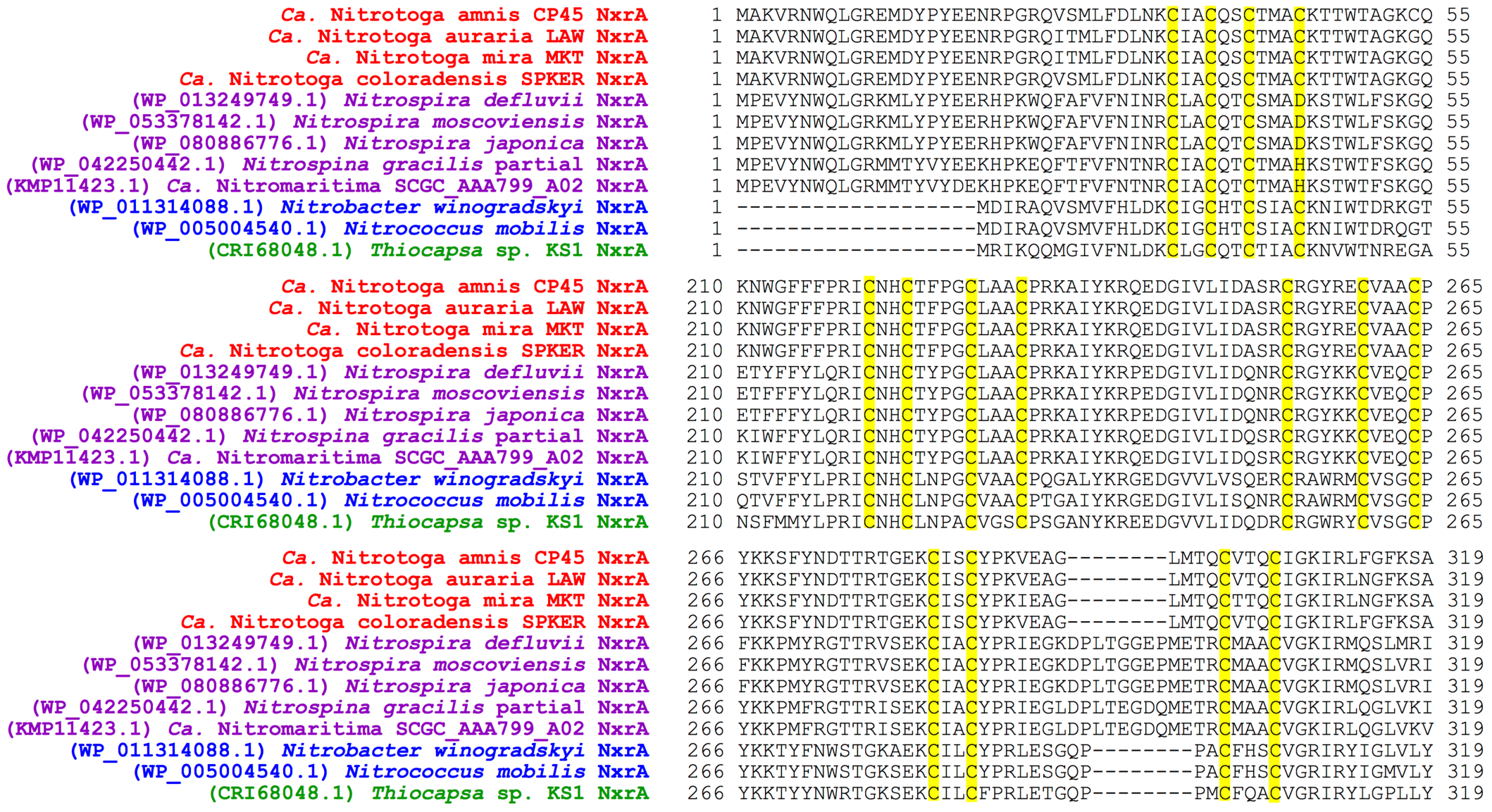
Partial alignment of important residues from selected NxrB subunits representing the *Nitrotoga* (Red), as well as canonical periplasmic-facing (Purple) and cytoplasmic-facing (Blue/Green) NXR. Highlights represent Fe-S cluster binding residues (Yellow).

**Supplemental Figure S3.**
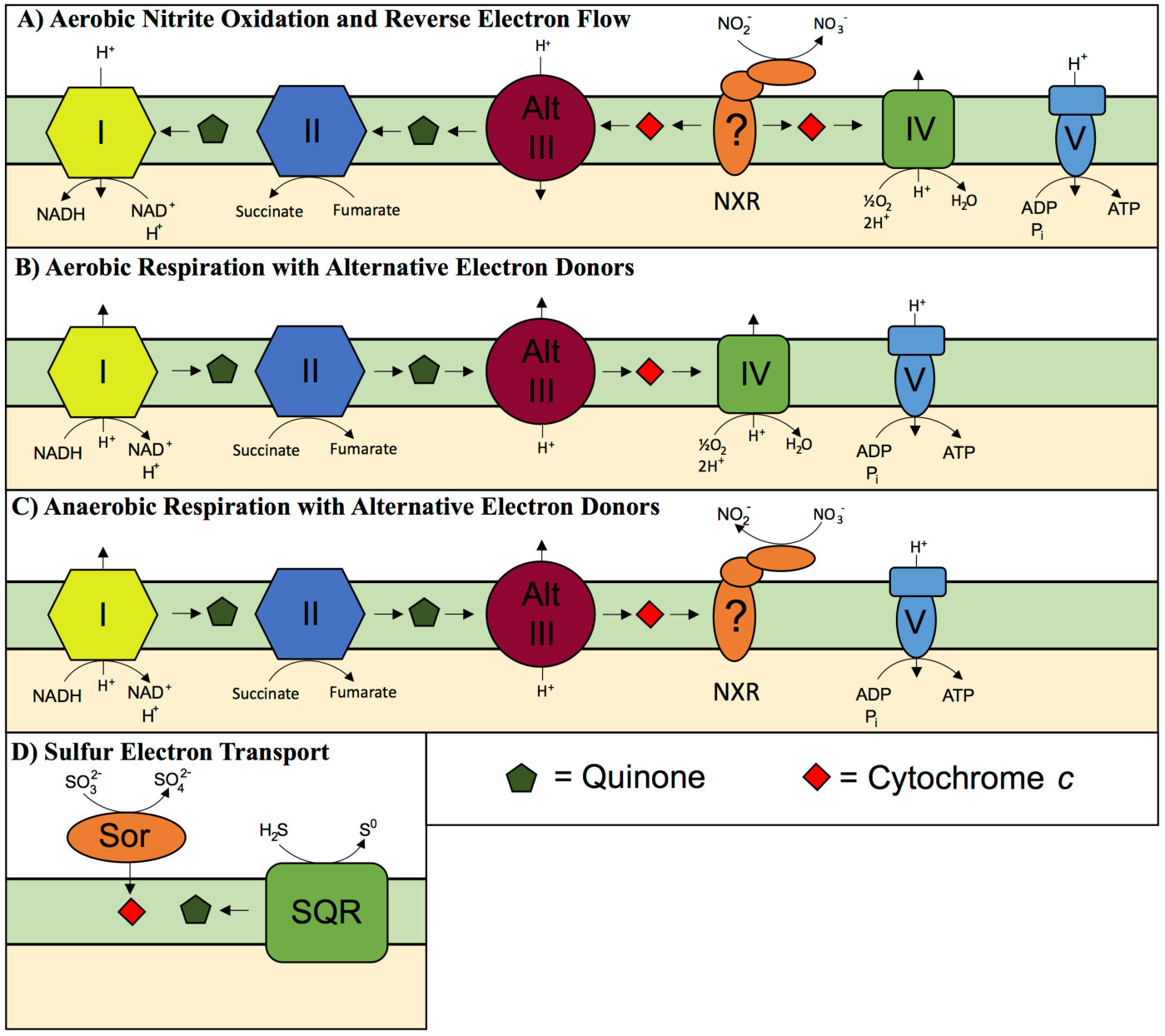
Schematic of electron transfer in *Nitrotoga* based on genomic evidence. **A)** Canonical nitrite oxidation performed by NXR will liberate two electrons onto cytochrome *c* and flow forwards to the terminal oxidase (Complex IV), or backwards to regenerate NADH via Alternative Complex III and the quinone pool, or to generate biochemical intermediates via Complex II. The question mark located in the NxrC subunit symbolizes the uncertainty in whether or not the holoenzyme is anchored to the cell membrane. **B)** Aerobic respiration with alternative electron donors such as NADH derived from organic carbon utilization. **C)** Hypothesized anaerobic respiration with alternative electron donors (e.g., NADH) with the reduction of nitrate to nitrite via NXR as seen for other NOB. **D)** Electrons derived from reduced sulfur compounds (sulfites or sulfides) are transferred to cytochrome *c* or quinone, which can enter at any point in electron transfer shown in panels B or C.

**Supplemental Figure S4.**
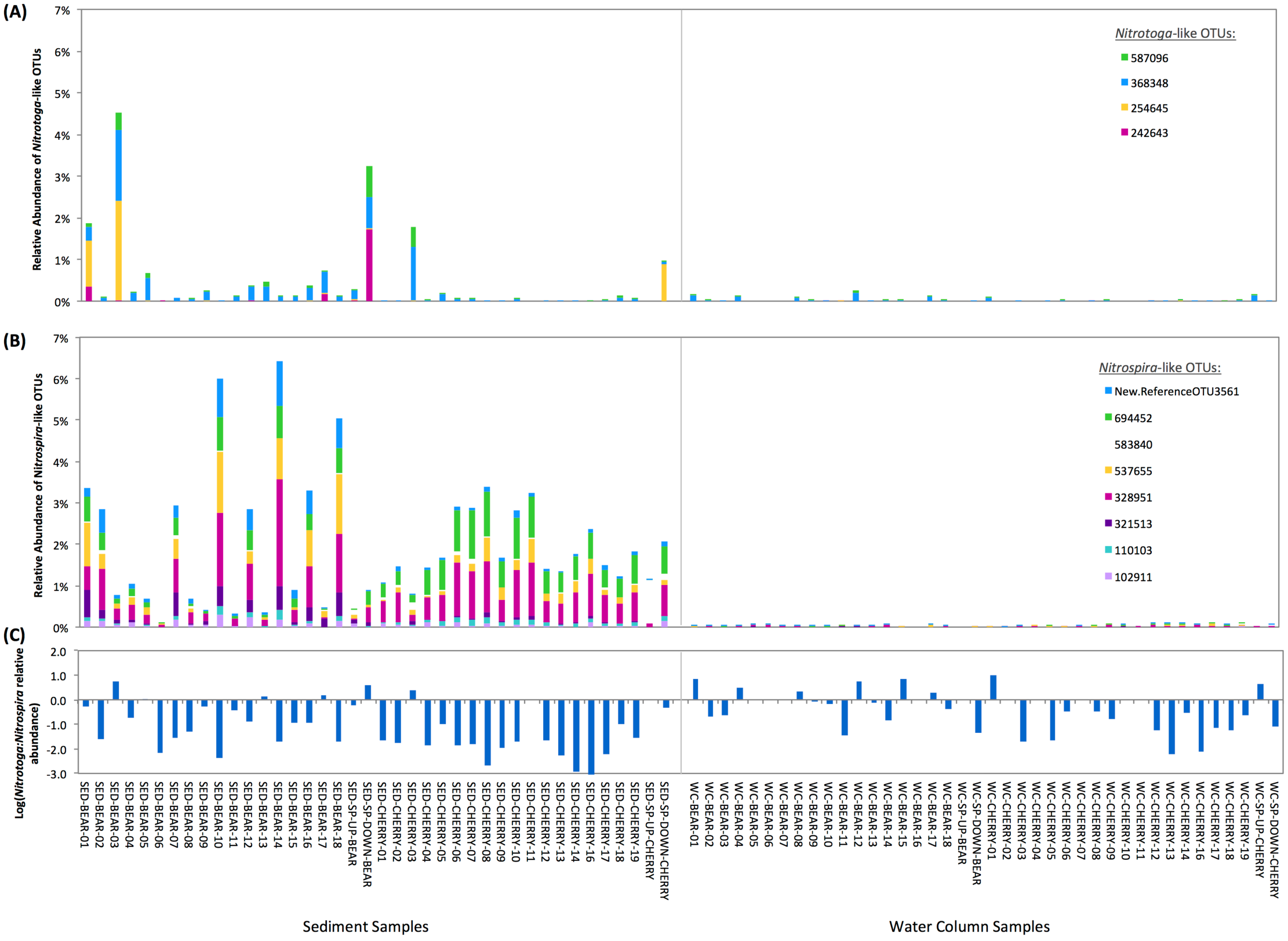
Relative abundance of **A)** *Nitrotoga-* and **B)** *Nitrospira*-like 16S rRNA gene sequence OTUs from water column (“WC”) and sediment (“SED”) samples in Bear Creek (“BEAR”), Cherry Creek (“CHERRY”), and the upstream (“SP-UP”) and downstream (“SP-DOWN”) sites at their respective confluences with the South Platte River. OTUs were based on the amplification with general 16S rRNA gene primers targeting the total bacterial community and were grouped at 97% nucleotide identity. *Nitrotoga* and *Nitrospira* OTUs were identified based on BLAST searches against the SILVA rRNA gene database. **C)** Log ratio of the summed relative abundance of *Nitrotoga*- to *Nitrospira*-like 16S rRNA gene sequences.

**Supplemental File 1.**
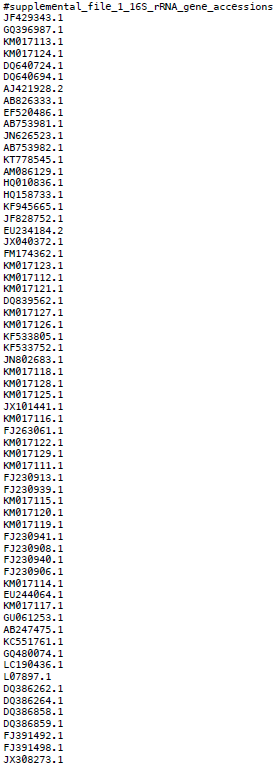
List of accessions for 16S rRNA genes used in Figure 3 and Supplemental Figure S1.

**Supplemental File 2.**
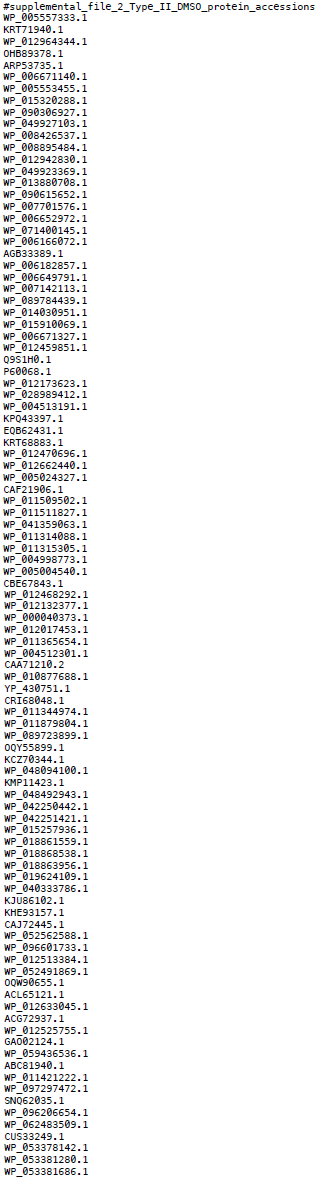
List of accessions for Type II DMSO reductase proteins used in Figure 4.

## SUPPLEMTENAL NOTE

### METHODS

#### Culture inoculation and growth

Freshwater Nitrite Oxidizer Medium (FNOM) was prepared by mixing 1 g NaCl, 0.4 g MgCl_2_-6H_2_O, 0.1 g CaCl_2_-2H_2_O, 0.5 g KCl, 100 μL 10X vitamin solution (Balch *et al*., 1979), 1 mL 1M NaHCO_3_, and 300 μL 1M NaNO_2_ per liter. The pH of the media was lowered to 7.0 using 10% HCl, then autoclaved. After autoclaving, 10 mL of separately autoclaved 4 g _*_ L^-1^ KH_2_PO_4_ and 1 mL trace metal solution (Biebl and Pfennig, 1978) were sterilely added to the media before storing at 4°C in the dark.

Two surface sediment samples were aseptically collected in February 2015 from the urban-impacted Cherry Creek in downtown Denver, CO (samples MKT and LAW) using a cut-off sterile 30 mL syringe and returned to the lab on ice. The same day, the top 0.5 cm of sediment was mixed with 10 mL sterile FNOM, then 1 mL of the sediment slurry was transferred to 100 mL FNOM for incubation at room temperature in the dark. Two water column samples were collected in May 2015 from two agriculturally-impacted rivers near Greeley, CO (about 100 km North of Denver, CO) (samples CP45 from the Cache La Poudre River and SPKER from the South Platte River). River water was kept on ice in the field and stored at 4°C upon return to the lab. After five days, 10 mL of each water sample was transferred to 100 mL FNOM and allowed to incubate at room temperature in the dark.

Nitrite consumption was regularly monitored in the cultures using a Griess nitrite color reagent (Griess-Romijn van Eck, 1966) composed of 0.5 g sulfanilamide, 0.05 g N-(1-naphthyl) ethylenediamine dihydrochloride, 5 mL 85% phosphoric acid, and MilliQ water to a final volume of 50 mL. Nitrite color reagent was mixed with the culture at a 1:10 ratio for visual estimates (+/-) or at a 1:1 ratio for quantitative spectrophotometric measurements.

To determine rates of nitrite oxidation, 100 μL of each sample line (CP45, LAW, MKT, SPKER) was inoculated into three bottles with 100 mL FNOM. At regular intervals, samples were collected in triplicate from each bottle and mixed with equal volumes of fresh Griess nitrite color reagent, then the optical density (OD) was measured at 540, 545, and 550 nm using a BioTek Synergy HT plate reader (BioTek, Winooski, VT). The mean maximum OD was used to calculate nitrite concentrations based on a standard curve of sterile media ranging from 0-0.3 mM nitrite with the Gen5 analysis software (BioTek, Winooski, VT). Sterile FNOM was used as a negative control. Nitrite oxidation rates were calculated across three time points (R^2^ value of 0.93-1.0) within logarithmic nitrite consumption for each bottle.

#### DNA extraction

At mid- to late-phase of exponential nitrite oxidation, 400 mL of each culture was filtered onto a 0.2 μm Supor 200 filter (Pall, New York, NY). Filters were cut into small pieces with a sterile scalpel and aseptically placed into a Lysing Matrix E Bead Beating Tube (MP Biomedicals, Santa Ana, CA) with 800 μL lysing buffer (750 mM sucrose, 20 mM EDTA, 400 mM NaCl, 50 mM Tris (pH 8.4)) and 100 μL 10% SDS. Samples were vortexed briefly before bead beating in a FastPrep-24 5G reciprocating homogenizer (MP Biomedicals, Santa Ana, CA) at 5 m/s for 30 seconds. Samples were incubated at 99°C for 1-3 minutes before adding 50 μL 20 mg/mL proteinase K, then incubated for 3-5.5 hours in a rotating hybridization oven at 55°C. Cold 100% ethanol (500 μL) was added to each sample in the same tube, and then DNA was purified using the DNeasy Blood and Tissue Kit (following manufacturer’s instructions for purification) (Qiagen, Hilden, Germany). Extracted DNA was quantified with a Qubit fluorometer (Thermo Fisher, Waltham, MA), using the High Sensitivity dsDNA assay.

#### Metagenome and *Nitrotoga* genome assembly

BBDuk v36.99 (http://jgi.doe.gov/data-and-tools/bbtools) was used to remove sequencing adapters and trim metagenomic reads (mink=8, hdist=1, qtrim=20, minlength=50, ftm=5, tpe, tbo). Read quality distributions were checked before and after trimming using FASTQC (https://www.bioinformatics.babraham.ac.uk/projects/fastqc/.) Metagenomes were assembled from each culture using the filtered and trimmed reads with MEGAHIT v1.0.6.1 (k-min=32, k-max=121, k-step=10). Reads were mapped to the metagenome assemblies using BBMap v36.x (http://jgi.doe.gov/data-and-tools/bbtools/) and contigs were binned using MetaBAT v.0.32.4 (-- verysensitive-B20 --unbinned) (Kang *et al.*, 2015). Preliminary taxonomy of each genomic bin was identified based on (1) BLAST searches of 16S rRNA gene sequences (assembled using EMIRGE (Miller *et al.*, 2011)) against the SILVA 16S rRNA gene database (release 123) (Quast *et al.*, 2013); and (2) the CheckM lineage workflow (Parks *et al.*, 2015). In each culture, only one putative NOB was identified belonging to the *Nitrotoga* genus.

Putative *Nitrotoga* bins of interest were manually refined using the Anvi’o metagenomics pipeline version 2.1.0 (Eren *et al.*, 2015). Putative *Nitrotoga* bins were combined in the CP45 and LAW metagenomes as these genomes were each split into two different bins. The CheckM merge function (Parks *et al.*, 2015) was used to supervise bin mergers and the lineage workflow was run again after merging to ensure the completeness estimates increased and contamination estimates did not. To check for contaminant and chimeric contigs, genes were called with Prodigal (Hyatt *et al.*, 2010) and BLASTP (Camacho *et al.*, 2009) was used to find the best hit for each predicted protein against the UniRef90 database (release 2016_11) (Suzek *et al.*, 2014). Contigs with suspicious BLASTP taxonomy results were further scrutinized and removed upon later reassembly as needed.

For an iterative reassembly process of *Nitrotoga* genomes, reads were mapped to contigs of individual bins using BBSplit v36.x (http://jgi.doe.gov/data-and-tools/bbtools/) and were assembled using SPAdes v3.9.0 (Bankevich *et al.*, 2012) under the “careful” setting with MEGAHIT-assembled contigs given as “trusted contigs” and the same kmer range as used in metagenome assembly. Assembly graphs were visualized using Bandage (Wick *et al.*, 2015) to identify suspicious contigs. Contigs were removed from the bin before further reassembly if they had dissimilar best UniRef90 taxonomy hits compared to the rest of the contigs, inconsistent BLASTX taxonomy hits against the NCBI nr database, and were not found to be present within bins of other *Nitrotoga* genomes (from this study).

16S rRNA gene sequences (which rarely bin properly in metagenomic assemblies) were manually added to the *Nitrotoga* genome assemblies. To avoid selection bias, all ‘unbinned’ MEGAHIT-assembled contigs were searched against the SILVA 16S rRNA database (release 128) (Quast *et al.*, 2013) using BLASTN (Camacho *et al.*, 2009). All contigs with an alignment ≥300 bp (≥189 for the LAW metagenome due to a *Nitrotoga* hit of that size) to any 16S rRNA gene from the database were added to the *Nitrotoga* genome assemblies. The resulting assembly graph was visualized in Bandage (Wick *et al.*, 2015), and the internal BLAST function was used to search for all added 16S rRNA suspected contigs. In each of the four *Nitrotoga* genomes, only one 16S rRNA gene assembled on a contig with paired reads mapped to other *Nitrotoga* genomic contigs. The assembled 16S rRNA genes from each culture were most similar to *Nitrotoga* sequences in SILVA. The correct contig containing the *Nitrotoga* 16S rRNA gene was kept, and others removed from the assembly.

A similar process was followed for adding contigs with nitrite oxidoreductase (*nxr*) gene sequences to the assembly. Predicted protein sequences from ‘unbinned’ MEGAHIT-assembled contigs were searched for members of the Type II DMSO reductase family (TIGR03479, TIGR03478, TIGR03477, TIGR03482) using HMMER3 (Eddy, 2011). All contigs with hits were added to the SPAdes reassembly if they had a coverage estimate that was similar to, or higher than, that of the respective *Nitrotoga* genome. Ultimately, a single unbinned contig in each of the CP45, LAW, and MKT metagenomes held all *nxr* genes. Eight unbinned contigs from the SPKER metagenome were used to assemble the *nxr* contig in the SPKER *Nitrotoga* genome.

Analysis of single nucleotide variants (SNVs) in Anvi’o (Eren *et al.*, 2015) indicated 32 SNVs across the SPKER *nxrA* gene, which are likely the result of variations between the three predicted *nxrA* copies in the SPKER *Nitrotoga* genome, and/or contaminant reads from one or more of the suspected *nxr* MEGAHIT-assembled contigs. Four of the SNVs were found in ~33% of mapped reads, indicating they may represent variants on one of the three suspected *nxrA* gene copies and caused contig breakage during assembly. None of the other *Nitrotoga nxrA* genes had SNVs and the SPKER *Nitrotoga nxrBCD* genes were intact with no SNVs. A consensus *nxrA* sequence is presented in this study for the SPKER *Nitrotoga* genome, as paired reads could not resolve the SNVs into individual gene copies.

#### Relative abundance of *Nitrotoga* in creek sediments and water column

Sediment samples from Bear Creek, Cherry Creek, and the South Platte River were collected using a sterile spatula and sterile petri dish. Prior to sampling at each site, the spatula was rinsed with 70% ethanol and wiped dry with a clean Kimwipe. The spatula was then rinsed with sterile water to remove residual ethanol, followed by site water. The bottom of the sterile petri dish was then placed down into the sediment, open-side-down. The sterile spatula was slid under the dish, trapping the sediment in the petri dish. The lid was put on the petri dish and wrapped in parafilm. All sediment samples were stored on dry ice for a maximum of three hours before arriving at the lab for permanent storage at −80°C.

Water samples were collected using sterile 1 L Nalgene bottles submerged approximately 15 cm in creek water undergoing constant flow. Nalgene bottles were rinsed with site water three times before sample acquisition. Following sample collection, samples were immediately placed on ice and transferred to the lab for processing for filtration within three hours. Water column samples were filtered onto 25 mm diameter, 0.22 μm pore size Supor membrane (Pall Corporation, Ann Arbor, MI) filters in a Swinnex (Merck Millipore, Burlington, MA) filter housing using a peristaltic pump (Series II Geopump, Geotech Environmental Equipment, Denver, CO). Following sample filtration, filters were stored at −80°C until DNA extraction. Pump tubing was sterilized prior to filtering each sample by sequential rinses of the pump tubing: (1) rinsed with 250 mL of autoclaved MilliQ water; (2) recirculation of 500 mL of 10% HCl through the tubing for three minutes; (3) rinsed again with 250 mL of autoclaved MilliQ water; and (4) rinsed with 250 mL of site water.

DNA was extracted using the MP Biomedicals FastDNA Spin Kit for Soil (MP Biomedicals, Santa Ana, CA) according to kit instructions with homogenization in the FastPrep-24 5G reciprocating homogenizer at 6.0 m/sec for 40 seconds. The Qubit dsDNA HS and dsDNA BR Assay kits (Life Technologies, Carlsbad, CA) were used to determine the DNA concentration of extracts. Qubit DNA quantitation was run in duplicate to determine the average DNA concentration for each DNA extract.

Sequence processing was conducted using QIIME (Caporaso *et al.*, 2010). Paired-end reads were joined using fastq-join and filtered to a Phred quality score of 20. Sequences with < 80 bp merge length were discarded. The last 20 bp was removed from both ends of sequences after merging to remove primers. Processed reads were clustered into operation taxonomic units (OTUs) at 97% sequence identity and the DECIPHER web tool was used to check for chimeras (short sequences) (Wright *et al.*, 2012). Putative chimeras as well as OTUs with relative abundance < 0.05% were removed. All chloroplast OTUs and OTUs found in sequencing controls were also removed before analysis. Final taxonomy was assigned using a BLAST search against the SILVA 16S rRNA database (release 128) (Quast *et al.*, 2013).

## RESULTS AND DISCUSSION

### *Nitrotoga* genome assembly

CheckM indicated that the *Nitrotoga* genomes were near-complete based on a collection of 419 single-copy gene markers conserved within the Betaproteobacteria (UID3959). The LAW, MKT, and SPKER genomes were predicted to be 98.2% complete, while the CP45 genome was 97.0% complete due to the loss of three markers that were present on small contigs (<2 kb) removed before annotation (PF00731.15 AIR carboxylase; PF01259.13 Phosphoribosylaminoimidazolesuccinocarboxamide synthase; and TIGR02392 alternative sigma factor RpoH). All genomes were missing the same five marker genes (TIGR01745 aspartate-semialdehyde dehydrogenase; TIGR01574 tRNA-i(6)A37 thiotransferase enzyme MiaB; PF03618.9 Kinase-PPPase; TIGR01161 phosphoribosylaminoimidazole carboxylase, ATPase subunit; and PF13603.1 Leucyl-tRNA synthetase, Domain 2). A manual search for each marker using HMMER3 (Eddy, 2011) revealed strong (evalue <1e-41) hits to four of the five missing genes in all genomes. The identification of these four markers improved completeness estimates to 99.8% complete. The fifth marker (PF03618.9) had no hits from the *Nitrotoga* predicted protein sequences and was not found on any contig with suitable coverage among the ‘unbinned’ contigs. Homologs to this marker gene are found in the closest sequenced relatives: *Sideroxydans lithotrophicus* ES-1, *Gallionella capsiferriformans* ES-2, and *Gallionella acididurans* ShG14-8.

*Nitrotoga* genomes were estimated to contain 0.24-0.3% contamination. Specifically, the CP45, LAW, MKT genomes had duplications of the same two markers (PF09976.4 Tetratricopeptide repeat-like domain; and PF08340.6 domain of unknown function 1732), while the SPKER genome only had a duplicate of PF08340.6. Secondary copies had a drastically reduced HMM hit (evalue 2-11 orders of magnitude higher), so they may represent false-positives. None of the published Gallionellaceae genomes showed duplications of these genes.

### *Nitrotoga* nitrogen metabolism

#### Nitrite oxidation

NxrA contains a specific nitrite/nitrate substrate binding channel, a molybdenum-bis(pyranopterin guanosine dinucleotide) (Mo-bisPGD) motif, and one iron-sulfur cluster for the conduction of electrons to the beta subunit (Daims, Lücker, & Wagner, 2016; Grimaldi, *et al*. 2013; Hille, Hall, & Basu, 2014; Magalon, *et al*. 2011). The beta subunit acts as an electron conductor, passing through three [4Fe-4S] clusters and one [3Fe-4S] cluster (Daims, Lücker, & Wagner, 2016; Grimaldi *et al*., 2013; Hille, Hall, & Basu, 2014; Magalon, *et al*. 2011). Finally, the variable gamma subunit, NxrC, is thought to bind 1-2 heme groups to transfer electrons to a cytochrome *c*, and is likely bound to the membrane, anchoring the NXR holoenzyme (Daims *et al.*, 2016). NOB genomes with periplasmic-facing NXR typically encode multiple candidate *nxrC* genes, with varying sizes and heme-binding components (Daims *et al*., 2015; Lücker *et al*., 2010; Lücker, *et al*., 2013; van Kessel *et al*., 2015)

*Nitrotoga nxr* genes were ultimately placed on single contigs within each genome forming an *nxrABCD* operon. *Nitrotoga* NxrA (TIGR03479, evalue < 2.2e-74) was 1169 amino acids in length in the CP45, MKT, and LAW genomes and 1155 aa in the SPKER genome due to a truncated N-terminus (Figure 4b). *Nitrotoga* NxrB (TIGR03478, evalue <2.6e-73) was 385 amino acids long in all genomes, while *Nitrotoga* NxrC (TIGR03477, evalue <2.8e-36) was 371 or 372 amino acids in length. Each contig also contained an NxrD delta subunit (TIGR03482, evalue <8.8e-26), which may act as a chaperone similar to TorD used in molybdenum cofactor assembly and protein folding (Hille *et al.*, 2014; Lücker *et al.*, 2013; Magalon *et al.*, 2011; Ngugi *et al.*, 2015).

Two conserved hypothetical proteins were found downstream of the *nxr* operon in most *Nitrotoga* genomes: a 70 amino acid protein of unknown function, and a 341 amino acid protein with conserved domains related to iron-transfer P-loop NTPases, which are required for cytosolic Fe-S cluster assembly (factor NBP35). These two genes are located on the same contig as the *nxr* genes in the CP45, LAW, and MKT genomes. The SPKER *nxr* contig contained only the *nxrABCD* operon, but the 341 amino acid protein was found on a neighboring contig. The CP45 *nxr* contig had additional truncated hypothetical proteins on either end of the contig.

*Nitrotoga* NXR subunits have a conserved protein structure that is highly divergent from other NXRs and members of the Type II DMSO reductase family of enzymes (Figure 4). *Nitrotoga* NxrA subunits were at least 98.1% identical to each other across the 1169 amino acid protein but were as low as 84.1% identical across the nucleotide alignment (SPKER vs LAW). Similar patterns were seen with the beta subunit (≥97.9% amino acid identity; ≥87.0% nucleotide identity), while the gamma subunit was slightly more divergent (≥92.8% amino acid identity; ≥85.3% nucleotide identity). The delta chaperone was also highly conserved between *Nitrotoga* genomes (≥96.4% amino acid identity; ≥87.6% nucleotide identity).

#### Dissimilatory and assimilatory nitrogen metabolism

Urea is an important source of ammonia for many NOB (e.g., *Nitrospina* and some *Nitrospira*) (Daims *et al.*, 2016; Ushiki *et al.*, 2018). A urea-binding-like protein was found in the LAW genome located near most of the nitrogen transport systems (i.e., *narK* nitrite/nitrate transporter, nitrogen metabolism transcription factor *ntrC*, formate/nitrite transporter, *nirK* assimilatory nitrite reductase, and ammonia transporter *amtB*). However, no urea transporter or urease genes were identified in any *Nitrotoga* genomes.

Each *Nitrotoga* genome contained a nitrilase enzyme that degrades nitriles (C≡N bonds) to ammonia and carboxylic acids. Nitrilase could potentially be used as defense against simple nitriles like cyanide or to cleave ammonia for assimilation within the *Nitrotoga* cells. Previously, a cyanate-degrading NOB (via a cyanase enzyme) was shown to participate in ‘reciprocal feeding’ by releasing ammonia for consumption by ammonia-oxidizers (Palatinszky *et al.*, 2015). However, no cyanate transporter or cyanase genes were identified in any *Nitrotoga* genomes. Each *Nitrotoga* genome did contain four unique genes with rhodanese domains, which are known to detoxify cyanide (CN^-^) using thiosulfate (S_2_O_3_^2-^) as a sulfur donor, producing thiocyanate (SCN^-^) and sulfite (SO_3_^2-^). One of these genes was predicted to be anchored into the cell membrane and face the periplasm. Thus, *Nitrotoga* may be capable of detoxifying cyanide while also contributing to their sulfur metabolism via sulfite production (see *sulfur energetics* section).

### *Nitrotoga* carbon metabolism

#### Carbon fixation

The CP45, LAW, and MKT genomes had two copies of both small and large ribulose 1,5-bisphosphate carboxylase (RuBisCO), the key enzyme for CO_2_ fixation, while the SPKER genome had single copies. All *Nitrotoga* genomes had an active form IC/ID RuBisCO, while the second copy in the CP45, LAW, and MKT genomes was related to form IA RuBisCO. All form I RuBisCO are likely active in carbon fixation (unlike most Form IV RuBisCO (Tabita *et al.*, 2007)) and subtypes IA, IC, and ID have been found across the Proteobacteria (Tabita *et al.*, 2008).

The two key enzymes of the reverse tricarboxylic acid (rTCA) cycle (oxoglutarate:ferredoxin oxidoreductase and ATP-citrate lyase) were missing from *Nitrotoga* genomes. The rTCA cycle is used in *Nitrospira*, *Nitrospina*, and *Ca.* Nitromaritima species to fix carbon dioxide (Lücker *et al.*, 2010, 2013; Ngugi *et al.*, 2015), while the *Nitrolancea*, *Nitrococcus*, and *Nitrobacter* utilize the Calvin cycle (Sorokin *et al.*, 2012; Füssel *et al.*, 2017; Starkenburg *et al.*, 2008).

#### Polysaccharide storage and Exopolysaccharide

Glycogen and starch storage is likely in *Nitrotoga* given the presence of key genes (i.e., glucose-1-phosphate adenylyltransferase, 1,4-alpha-glucan branching enzymes, alpha-amylase, starch synthase, and starch phosphorylase). A pathway for cellulose production (cellulose synthase (UDP-forming)) and hydrolysis (cellulase/endogluconase and cellobiose phosphorylase) to glucose 1-phosphate was also present in *Nitrotoga* genomes, except the LAW genome was missing a cellobiose phosphorylase gene.

Exopolysaccharide (EPS) synthesis is also likely in *Nitrotoga*, as an extensive array of PEP-CTERM exopolysaccharide sorting enzymes (Haft *et al.*, 2006), polysaccharide export pumps, and saccharide modifiers (i.e., UDP-glucose dehydrogenase, polyprenyl glycosylphosphotransferase, D-alanyl-lipoteichoic acid acyltransferase) were present in all genomes. EPS production has been observed in *Nitrotoga* via microscopy (Ishii *et al.*, 2017), and may explain the formation of microcolonies in WWTPs (Lücker *et al.*, 2015).

### *Nitrotoga* iron acquisition

The CP45, LAW, and SPKER genomes included two genes with sequence similarity to IucA/IucC and FhuF domains that are commonly found in nonribosomal peptide synthase-independent siderophore biosynthesis (Challis, 2005). However, it is unclear whether *Nitrotoga* are capable of complete siderophore biosynthesis as these genes also resemble a ferric reductase, which may be used to reduce siderophore-bound iron prior to transport across the cell membrane. Reduced Fe^2+^ is likely transported into the cytoplasm via an EfeU Iron/Lead type transporter, while an FhuDBC ABC-type transporter can transport some complete siderophores across the cell membrane. An AfuABC ABC-type transporter was also found with predicted Fe^3+^ transporter activity, however recent evidence suggests this may actually be a phosphorylated carbohydrate pump (Sit *et al.*, 2015).

Fe^3+^ storage is possible with bacterioferritin (BfrB) and a bacterioferritin-associated ferredoxin (Bfd) encoded in *Nitrotoga* genomes. However, a ferredoxin NADP reductase (Fpr) responsible for transferring electrons to Bfd, and ultimately Fe^3+^ for mobilization into the cytoplasm (Rivera, 2017; Wang *et al.*, 2015), was not detected in *Nitrotoga* genomes. A short ferritin-like protein, also useful for iron storage, was encoded in the CP45, LAW, and SPKER genomes although its function is unknown. Fur family transcriptional regulators, which measure cytoplasmic iron concentrations, were found near the *afuABC* operon and a zinc/cadmium transporter in *Nitrotoga* genomes and may respond to changes in iron availability.

A heme-degrading monooxygenase gene, *hmoA*, used to harvest heme from a host, was found in all *Nitrotoga* genomes. The presence of two putative hemolysin genes in all *Nitrotoga* genomes, and an additional heme oxygenase in the CP45 genome, may support an antagonistic role of *Nitrotoga* in bacterial communities. However, heme binding and uptake systems were not observed, and only the LAW genome encodes a Type I protein secretion system.

